# Ecological success of sexual and asexual reproductive strategies invading an environmentally unstable habitat

**DOI:** 10.1101/2020.02.10.942466

**Authors:** Willian T.A.F. Silva, Anna Nyqvist, Per R. Jonsson, Karin C. Harding

**Affiliations:** Department of Biological and Environmental Sciences, University of Gothenburg, Medicinaregatan 18, SE-413 90 Gothenburg, Sweden; Gothenburg Global Biodiversity Centre, Gothenburg, Sweden; Department of Marine Sciences, Tjärnö Marine Laboratory, University of Gothenburg, SE-452 96 Strömstad, Sweden

**Keywords:** metapopulation, geographic parthenogenesis, parthenogenesis, automictic, apomictic, invasion, population dynamics

## Abstract

Many aspects of sexual and asexual reproduction have been studied empirically and theoretically. The differences between sexual and asexual reproduction within a species often lead to a biased geographical distribution of individuals with different reproductive strategies. While sexuals are more abundant in the core habitat, asexuals are often found in marginal habitats along the edge of the species distribution. This pattern, called geographic parthenogenesis, has been observed in many species but the mechanisms reponsible for generating it are poorly known. We used a quantitative approach using a metapopulation model to explore the ecological processes that can lead to geographic parthenogenesis and the invasion of new habitats by different reproductive strategies. We analyzed the Allee effect on sexual populations and the population sensitivity to environmental stress during the invasion of a marginal, unstable habitat to demonstrate that a complex interaction between the Allee effect, sensitivity to environmental stress and the environmental conditions can determine the relative success of competing reproductive strategies during the initial invasion and longterm establishment in the marginal habitat. We discuss our results in the light of previous empirical and theoretical studies.

**Author Summary:** Individuals can reproduce with or without sex. Very often, closely related species are distributed in a such a way that the sexually reproducing species is most frequently found in the core habitat while the asexually reproducing species is found on the edge of the habitat range. This biased distribution of reproductive strategies across a habitat range is called geographic parthenogenesis and has been observed in several species. While many processes have been proposed to explain such a pattern, a quantitative approach of the ecological processes was absent. We investigated important differences between sexual and asexual reproduction and how these differences affect the success of sexuals and asexuals invading a marginal, unstable environment. We showed that the relative frequency of each reproductive strategy in the marginal habitat depends on how much sexuals rely on population density to reproduce and how much asexuals are affected by environmental stress relative to sexuals. Our study presents a quantitative ecological explanation for geographic parthenogenesis and provides the conditions under which different distribution patterns can emerge.

## Introduction

Niche and habitat range expansion are key ecological processes in both population dynamics and interspecific competition. These processes are essentially dependent on the efficiency of the reproductive strategies that are characteristic of each population in the new niches or habitats. Therefore, understanding how different reproductive strategies can affect these processes is fundamental for understanding population dynamics and interspecific competition during invasion of previously uninhabited habitats [1]. Although many different reproductive strategies exist, they can all be defined based on the occurrence of fusion of gametes provided by different individuals (sexual) or absence of such process (asexual). These different strategies have different benefits and disadvantages; for example, while asexual reproduction gives individuals the independence to reproduce without the need to find compatible mating partners [2; 3], sexual reproduction generates genetic diversity that may keep the population alive under adverse environmental conditions [4; 5; 6; 7]. Additionally, in many species, these reproductive strategies are not mutually exclusive, with individuals frequently switching between sexual and asexual reproduction [facultative modes of reproduction; 8; 9; 10; 11] or different populations having different reproductive strategies.

Because of the differences between sexual and asexual reproduction within a species, the geographical distribution of individuals with different reproductive strategies often differs [12; 13]. While sexuals are more abundant in the core habitat, where the species might have existed for a longer time, asexuals are often found in marginal habitats along the edge of the species distribution [14; 15; 16; 17; 18]. This pattern can be observed along the south-north gradient in the Northern Hemisphere, where glaciers have repeatedly wiped out northern populations and left the landscape open for recolonization from the south [19; 20], and also along elevation gradients in mountains, where climate becomes gradually harsher with altitude [21]. Similar patterns have been observed in aquatic environments along the salinity and temperature gradients, for example, in river-estuary complexes [22] and in the Baltic Sea [23; 24; 25; 26], where gradient extremes may be physiologically stressful for certain reproductive strategies. The mechanisms behind this spatial segregation (geographic parthenogenesis) are hotly debated and have been attributed to both evolutionary [e.g., 18; 19] and neutral random processes [27].

Genetic distribution patterns of geographic parthenogenesis have been investigated with empirical and theoretical approaches, both in plants [15; 28] and animals [14; 21; 29]. For example, in the alpine plant species *Ranunculus kuepferi*, apomictic (asexual) populations exhibit high genetic admixture near sexual populations but are highly uniform in remote areas, with few well-supported genetic clusters [30], indicating the occurrence of multiple colonization events by genetically different founders. However, in the genera *Taraxacum* (dandelion) and *Chondrilla* (skeleton weed), apomictic populations exhibit high genetic diversity, which can be explained by crosses between apomictics and sexuals (in regions where these reproductive strategies are sympatric) followed by colonization of marginal regions, or crosses between facultative apomictics in purely apomictic regions [31; 32]. In *Daphnia pulex* (water flea), asexual populations also exhibit elevated individual heterozygosities introduced by outcrosses [33], allowing outcrossed asexuals to displace sexuals due to the competitive advantages confered by their admixed genotypes. Although these genetic differences between sexual and asexual *Daphnia* can explain the differences in the geographic distribution of different reproductive strategies, it is also possible that genetic differences can cause ecological differentiation between reproductive strategies, allowing them to coexist in the same habitat, as has been suggested by experiments [34]. In many cases, however, genetic diversity in parthenogenetic populations is generally low [35; 36; 37; 38; 39], which may be explained by the invasion of marginal habitats by a small number of asexual individuals.

Several theoretical aspects of geographic parthenogenesis have been studied [1]. The evolution of spatial segregation between sexual and asexual populations was explored in annual hermaphrodites with an individual-based model [20]. In the model, the metapopulation consisted of patches arranged along a south-north axis, with reproductive rate gradually decreasing in the north direction and each patching favoring a locally adapted phenotype. Population dynamics led to asexual individuals exhibiting higher frequencies than sexual individuals in the north patches, which was explained by the gene flow from the south constraining sexual individuals to evolve local adaptations to the habitats in the north, while asexuals maintained their locally advantageous genotype once it had appeared. It has been suggested that asexuals that are well adapted to marginal habitats retain their adaptation while sexuals can suffer from gene flow of suboptimal alleles from the core-habitat. Additionally, populations with metapopulation dynamics tend to show geographic parthenogenesis because of the higher tolerance of asexuals to population bottlenecks and drift, allowing them to invade marginal habitats in small numbers [16]. On shorter time scales asexuals may also be better colonizers because they can reproduce without mating, thus avoiding the Allee effect that sexuals are subject to [16; 18]. A different model explored the effects of sexual conflict and mate limitation on the frequency of facultative parthenogens [40]. The magnitude of these effects together with levels of environmental productivity were primary determinants of the spatial distribution of different reproductive strategies (sexual or facultative parthenogenesis). Parthenogenesis was particularly favored when low environmental productivity caused low population density at the edges of the habitat, making mating either too difficult or too costly. Other models have also resulted in distribution patterns of reproduction strategies that is typical of geographic parthenogenesis [e.g., 41].

However, many important aspects of geographic parthenogenesis remain, surprisingly, unexplored. The existence of different types of asexual reproduction [42; 43] and the ability to change between different reproductive strategies are important factors that need to be addressed. The magnitude of the Allee effect, which can be particularly important in sexual populations [44; 45; 46], may play an important role in leading to geographic parthenogenesis but has been essentially ignored. And finally, population sensitivity to environmental variation in marginal habitats may determine whether habitat range expansion is possible and which reproductive strategies can be the most successful during range expansion, and yet these factors have been overlooked in particular from the empirical perspective. Because of the necessity to address these processes as leading causes of geographic parthenogenesis, we developed a metapopulation model of competition between different reproductive strategies and assessed the ecological success of each competing strategy during short-term invasions and long-term establishment in unstable habitats, under different environmental conditions. Our model considers both the Allee effect (weak and strong) that affects sexually reproducing populations and the population sensitivity to stressful environmental conditions, which can be affected by genetic diversity. We use the temporal mean population size during initial invasion and long-term establishment as a measure of the ecological success of each competing strategy.

## Methods

We model population dynamics in a metapopulation consisting of two qualitatively different patches. One patch (South patch) is assumed to host the larger, ancestral population, while the other patch (North patch or marginal habitat) is assumed to be a habitat that has been recently made available to the species under focus. The availability of a patch to a certain species can be affected by several factors (e.g., global warming may make northern ecosystems available to species that are typical from temperate climates). In our model, the South and North patches have different environmental properties, which are independent from the properties of the populations that they host. The South patch is characterized by a high environmental stability and hosts an ancestral population in equilibrium (Figure 1). Although, we use only two patches in the current study, the equations can be adapted to any number of patches.

**Figure 1:**
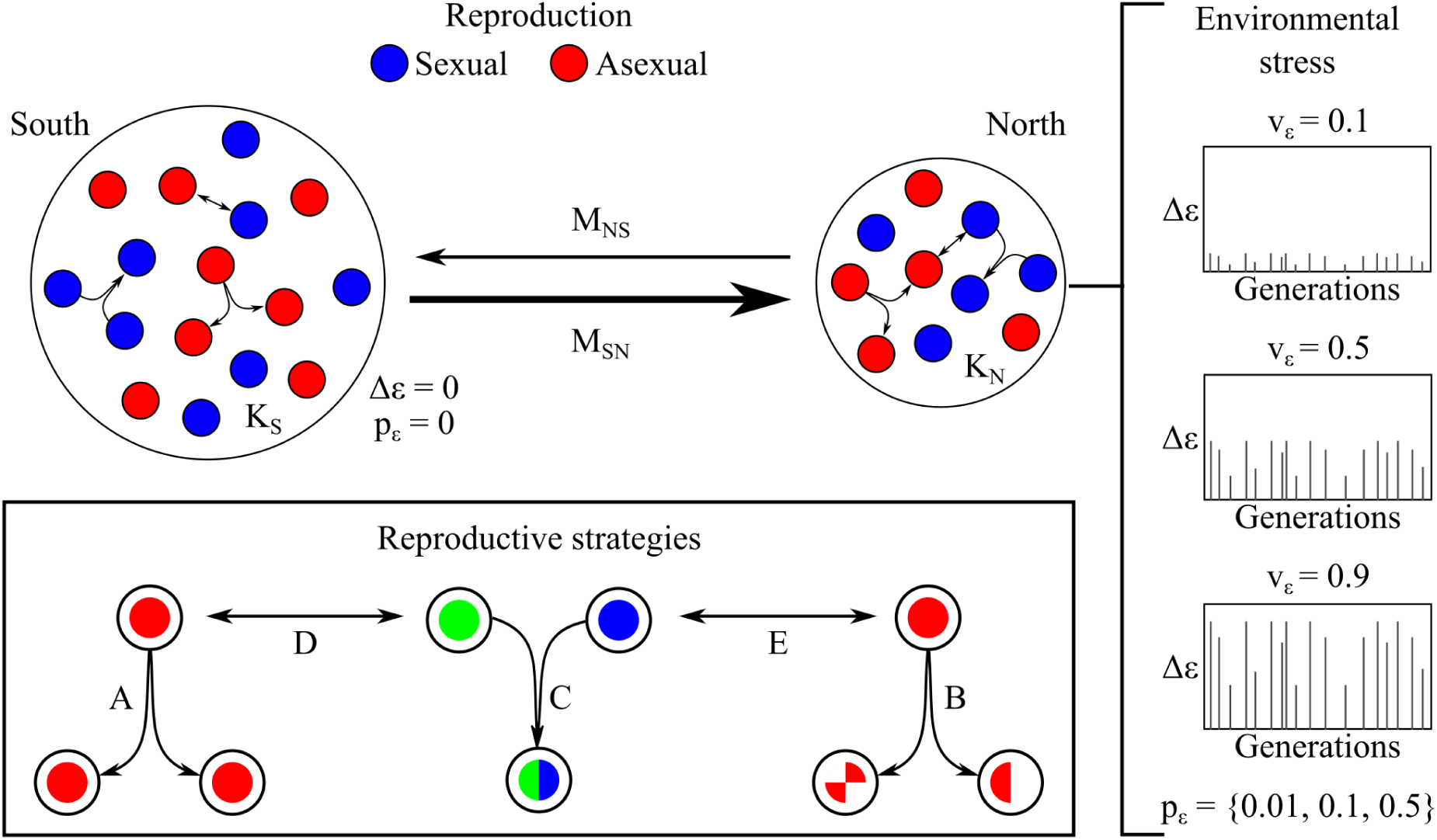
Graphical overview of the model design. Population dynamics include strategy-specific growth, transition and migration between an ancestral population (South patch) and a marginal habitat (North patch) where environmental conditions are unstable relative to the South patch. The panels to the right represent examples of three environmental regimes to which the North patch is exposed. Five different reproductive strategies were considered (bottom rectangle): obligate apomictic parthenogenesis or clonal reproduction (strategy A), obligate automictic parthenogenesis (strategy B), obligate sexual reproduction (strategy C), facultative apomictic parthenogenesis (strategy D) and facultative automictic parthenogenesis (strategy E). Complete description of the reproductive strategies in the main text. The subscripts/superscripts *S* and *N* indicate variables characterizing the South and North patches, respectively. The South patch differs from the North patch in its carrying capacity (*K*_*S*_ > *K*_*N*_), probability of occurrence of stressful events (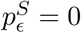 and 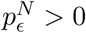), maximum environmental stress level (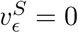 and 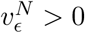) and effective stress level (Δ*ϵ*^*S*^ = 0 and Δ*ϵ*^*S*^ ≥ 0). *M* represents migration between patches.

The ancestral population consists of individuals that reproduce according to one of the following strategies, characterized by the properties of their growth rate:

A. Obligate apomictic parthenogenesis (clonal reproduction): Individuals do not need to find a mating partner in order to reproduce, which can be advantageous when population density is low. However, parents and offspring are genetically identical (very limited genetic variation), which makes the population very sensitive to changes in environmental conditions.
B. Obligate automictic parthenogenesis (non-clonal asexual reproduction): Individuals do not need to find a mating partner in order to reproduce, but unlike strategy A, parents and offspring are genetically different (to some extent) due to recombination during gamete production (limited but existent genetic variation). Because of the higher genetic variation, strategy B is assumed to be less sensitive to changes in environmental conditions than strategy A.
C. Obligate sexual reproduction: Individuals need to find a compatible partner to mate with and parents and offspring are genetically different. Because of the difficulty in finding compatible mating partners when the population density is low, strategy C is affected by the Allee effect. With sexual reproduction, however, the population keeps a higher genetic diversity that can be beneficial when environmental conditions change.
D. Facultative apomictic parthenogenesis: Individuals can transition between strategies A and C after assessment of population density. Strategy D individuals are affected by the same factors that affect strategies A and C when individuals act like such strategies. Strategy D individuals reproducing asexually are designated D- and individuals reproducing sexually are designated D+.
E. Facultative automictic parthenogenesis: Individuals can transition between strategies B and C after assessment of population density. Strategy E individuals are affected by the same factors that affect strategies B and C when individuals act like such strategies. Strategy E individuals reproducing asexually are designated E- and individuals reproducing sexually are designated E+.

It might be argued that strategy B (automictic parthenogenesis) is more sensitive to environmental stress than strategy A (apomictic parthenogenesis) because of the loss of heterozygosity in strategy B during reproduction. However, we consider the case where genomic diversity produced by automictic parthenogenetic reproduction is more beneficial than genomic homogeneity under natural selection caused by environmental stress because homozygosity caused by automictic parthenogenesis introduces new phenotypes into the population.

Different strategies can affect one another indirectly through their effects on the total population density and transitions between reproductive strategies (sexual to asexual and vice-versa), as explained below. For ease of reference, we define subpopulation as the fraction of the patch-specific total population that is composed of individuals with a specific reproductive strategy.

### Population dynamics

Population dynamics follow a logistic population growth model (Ricker model) with a variable growth rate, which is affected by environmental quality (carrying capacity), the Allee effect on strategies C-E, and environmental effects. Additionally, the total population density is affected by migration from/to the South patch. Change in subpopulation size from generation *t* to generation *t* + 1 due to growth is defined by the following difference equation:

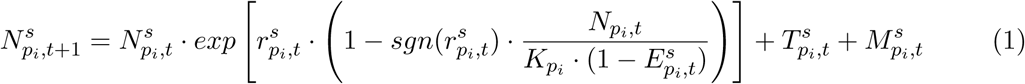

where the exponential term determines the logistic population growth rate based on the Ricker model and is equivalent to *λ* in exponential growth models, 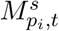 is the net change in population size due to migration South-North or vice-versa and 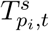 is the number of transitions from sexual to asexual reproduction and vice-versa in strategies D-E. In strategies A-C, 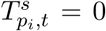. It is important to note that transitions follow growth and migration follows transitions, which means that transitions are calculated based on the outcome of the logistic term and migration is calculated after transitions happen. In the logistic growth term, 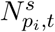 is the subpopulation size of the reproductive strategy indicated by the superscript *s* in the patch indicated by the subscript *p*_*i*_ (*p*_*S*_ for patch in the South and *p*_*N*_ for patch in the North) at time *t* (given in generations), 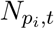 is the patch-specific total population size, 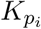 is the patch-specific constant maximum carrying capacity and 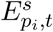 is the effect of changes in environmental conditions (explained in detail below). The variable 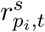 is the effective growth rate for the species under analysis after the reduction due to the Allee effect in strategies C-E. In strategies A and B, 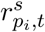 is simply the intrinsic maximum growth rate (explained below). In order to account for possible negative effects caused by both the Allee effect and reduction in environmental quality, it is necessary to introduce 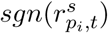 so that multiple negative effects do not cancel each other.

### Growth rates

All reproductive strategies have an equal intrinsic maximum growth rate *r*_*max*_, which leads to exponential growth when the following conditions are met: unlimited resources or maximum environmental quality 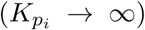, no Allee effect 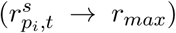, no environmental effect 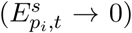 and no migration 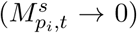. The maximum growth rate represents a biological limit in reproduction in the species under focus. By setting *r*_*max*_ = 1.1, we limit reproduction such that each individual can produce on average at most *λ* = *e*^1.1^ ≈ 3 offspring per generation (Figure S1).

### Allee effect

The Allee effect is present in the sexually reproducing subpopulations (strategies C-E) and accounts for the difficulty in finding compatible mating partners when the population size is small. In strategies D-E, the Allee effect only affects sexuals (D+ and E+). In asexual individuals (strategies A, B, D- and E-), 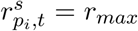 because mating is not necessary in order to reproduce.

The Allee-dependent growth rate 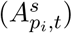 is lowest (*r*_*min*_) when the number of sexuals 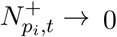 and increases as 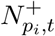 increases, until it reaches a biological limit (*r*_*max*_). This dynamic growth rate is defined by the following equation:

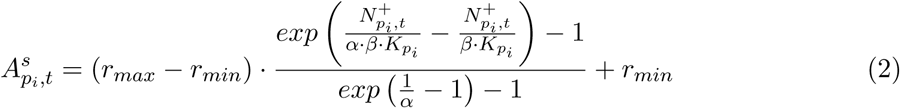

where *α* is the curvature of the function and 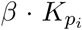 is the population size (relative to the carrying capacity 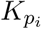) at which 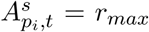, that is, when growth rate reaches its biological limit (Allee saturation point), such that 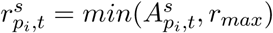 (Figure 2).

**Figure 2:**
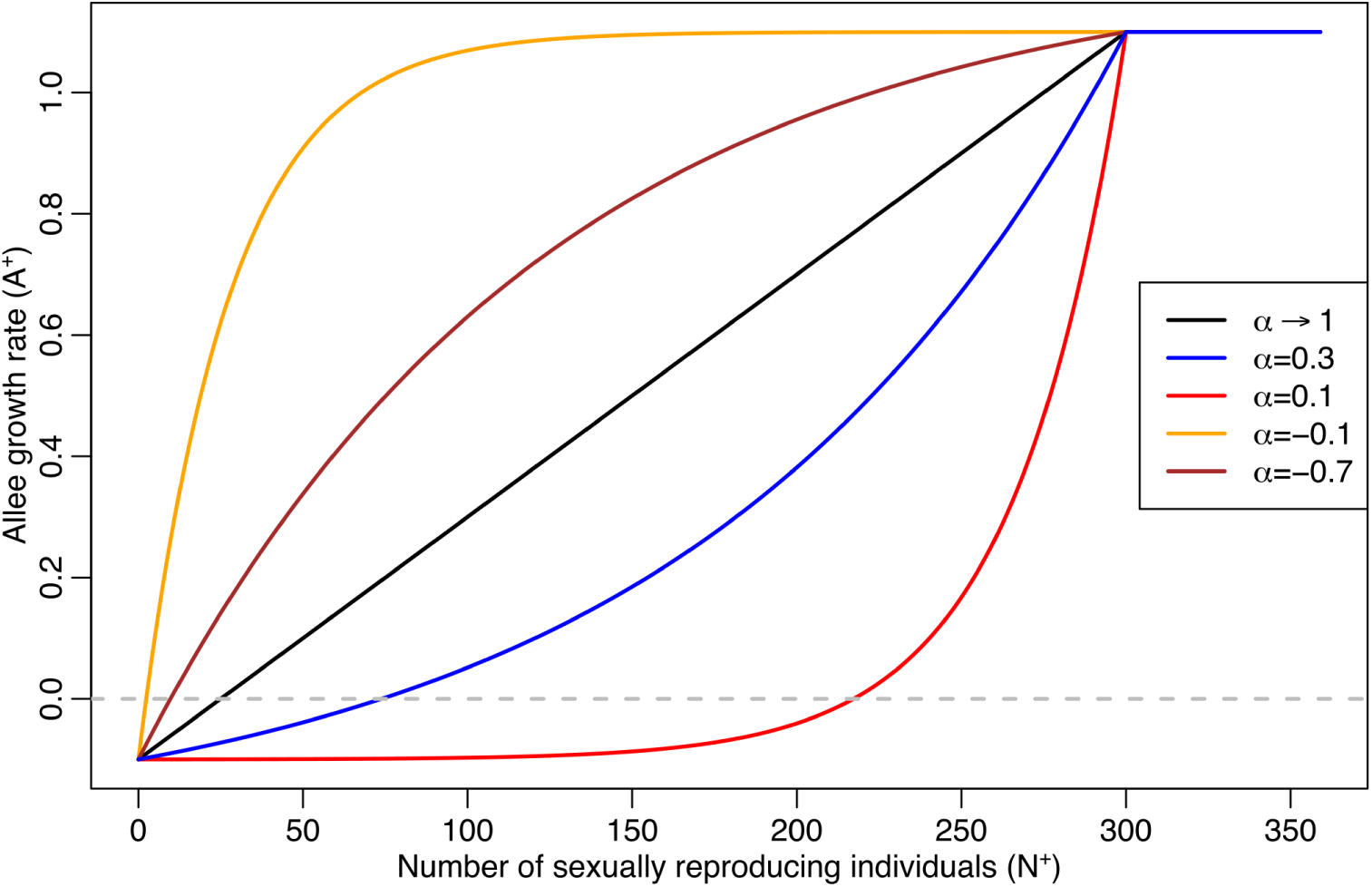
Allee function for different values of *α*. Parameter values used: *r*_*min*_ = −0.1, *r*_*max*_ = 1.1, *β* = 0.3, 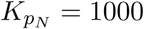

Unlike previous models of the Allee effect, we assume that growth rate increases with population density (with a monotonic relationship) and it is bound by an explicit upper biological limit. Difficulty in finding a compatible mating partner should always decrease with an increasing population density and Equation 2 allows us to explore this effect by changing the rate of change in growth rate (changes in *α* and *β*) as population density increases.

Our Allee model of growth rate has the following properties: (i) 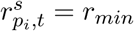 when 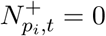; (ii) 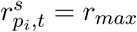 when 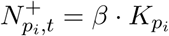; and (iii) 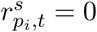 (no growth) when

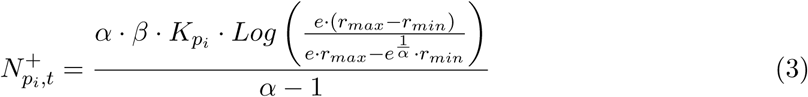

We set *r*_*min*_ = −0.1 (such that *λ* = *e*^−0.1^ ≈ 0.9) and assume that a very small subpopulation size (below the Allee threshold indicated by Equation 3) of sexuals results in many individuals dying before they have the chance to reproduce (negative net growth). Furthermore, we set 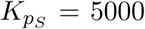 and 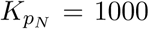 and explore values of *β* ∈ {0.1, 0.2, 0.3}, which means that the maximum growth rate of the sexually reproducing subpopulation in the North patch is achieved when the subpopulation size reaches 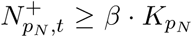 (which corresponds to 100, 200 and 300 individuals, respectively, as the population size at Allee saturation). The shape of the curve of the Allee effect was explored by setting *α* ∈ {−0.7, 0.1, 0.3, ∼ 1.0} in different simulations. When environmental conditions do not have an active effect on population dynamics 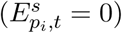, population growth is determined by the Allee effect and the carrying capacity of the patch, reaching its maximum when 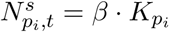. In weak Allee effect scenarios, we set *α* = 0.3 and *β* = 0.1; in strong Allee effect scenarios, we set *α* = 0.1 and *β* = 0.3 (Figure S2).

### Environmental conditions

Environmental stress can decrease the population growth rate because the population is not adapted to the new environmental conditions (e.g., starvation, diseases, energetic stress, reduction in resource availability due to anthropogenic disturbances). In our model, environmental stress reduces the effective carrying capacity of each reproductive strategy in the North patch, which is defined by 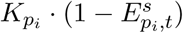 in Equation 1.

Because we are interested in how environmental stress affects population dynamics, we directly modeled environmental stress levels (Δ*ϵ*_*t*_) as scaled deviations from optimal environmental conditions 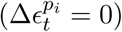 rather than the environmental variable itself. Our model assumes that environmental stress levels 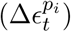 have the effect of reducing the patch-specific carrying capacity and therefore has a range 0 ≤ Δ*ϵ* ≤ 1, with Δ*ϵ* = 0 representing the complete absence of stress (optimal conditions) and Δ*ϵ* = 1 representing maximum stress, with complete reduction of the effective carrying capacity. Environmental stress occurs according to the following equation:

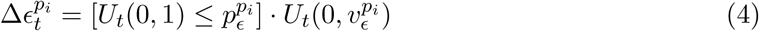

where 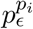 is the patch-specific probability of deviation from optimal environmental conditions and 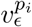 is the patch-specific maximum deviation (maximum stress level). The term 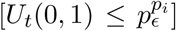 is a boolean indicator of presence (1) or absence (0) of change. Stressful environmental events across generations are stochastic and take a value drawn from a uniform distribution 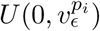. We explored the effects of different probability values 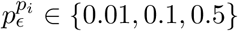 and maximum stress levels 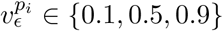 (Figure S3) on the initial invasion and long-term establishment of different reproductive strategies in the marginal habitat (North patch).

Although environmental stress affects the effective carrying capacity of the patch, different reproductive strategies are sensitive to these changes at different degrees due to their different genetic/phenotypic variation. We thus define 0 ≤ *ϕ*^*s*^ ≤ 1 as the sensitivity to changes in the effective carrying capacity of a patch due to environmental stress. We explored how sensitivity can affect the ecological success of each strategy. Unless indicated otherwise, sensitivity to environmental stress was assumed to be high in strategy A (clonal reproduction; *ϕ*^*A*^ = 1.0, used as a reference), intermediate in strategy B (non-clonal asexual reproduction; *ϕ*^*B*^ = 0.75) and low in strategy C (sexual reproduction; *ϕ*^*C*^ = 0.5) as a consequence of their genetic/phenotypic diversity. Note that strategy B differs from strategy A only in its sensitivity (*ϕ*^*B*^ < *ϕ*^*A*^). This sensitivity effect is assumed to be a linear function of the environmental stress level, such that:

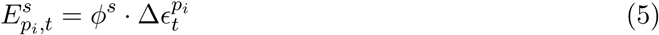

Equation 5 indicates that environmental effects on population dynamics are absent when environmental conditions are optimal 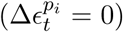 or the population is insensitive to changes in the environmental factors under focus (*ϕ*^*s*^ = 0). As noted earlier, strategies D and E share the properties of strategies A/C and B/C, respectively.

### Reproductive transitions

In facultative parthenogenetic strategies (D and E), transitions between sexual and asexual reproduction (and vice-versa) happen at a maximum rate *τ* and are affected by population density. We assume that individuals cannot distinguish between those that are reproducing asexually and those that are potential mating partners (sexual reproduction), so population assessment is based on total population size. After assessment of population density, a proportion of strategy D individuals transition from D- (asexual) to D+ (sexual) and vice-versa. Similarly, a proportion of strategy E individuals transition from E- (asexual) to E+ (sexual) and vice-versa. The net number of transitions from asexual (D-/E-) to asexual (D+/E+) reproduction 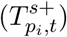 is calculated by the following equation:

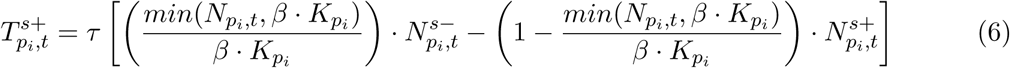

Equivalently, the net number of transitions from sexual (D+/E+) to asexual (D-/E-) reproduction 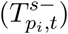 is calculated by the following equation:

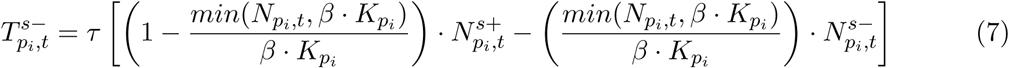

where 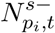 is the patch-specific number of individuals in strategy D/E reproducing asexually (D-/E-) and 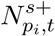 is the patch-specific number of individuals in strategy D/E reproducing sexually (D+/E+).

It is important to note that we assume no cost associated with the ability to transition in strategy D/E and that these strategies are only reproducing sexually in the ancestral population because population size is above the Allee saturation. We assumed *τ* = 0.2 across all simulations.

### Migration

Migration from/to the ancestral population in the South patch is density-dependent and is assumed to be greater in the north direction to stress the importance of habitat expansion in our model. Net migration (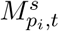, in number of individuals) is defined by the following equation:

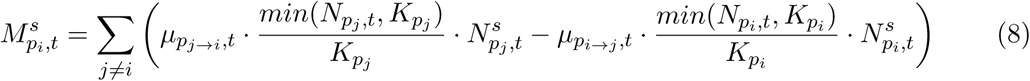

where 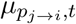 is the effective migration rate from patch *p*_*j*_ to patch *p*_*i*_ and 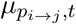 is the effective migration rate from patch *p*_*i*_ to patch *p*_*j*_.

We made migration stochastic by defining the effective migration rate according to the following equation:

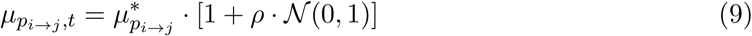

where 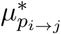 is the mean migration rate, *ρ* is the magnitude of the stochasticity and 𝒩(0, 1) is a standard normal deviate. We assumed 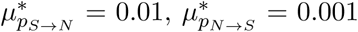 and *ρ* = 0.1 across all simulations.

### Ecological success during pairwise competition between strategies

Due to the stochastic environmental changes affecting population dynamics throughout time, the population structure at the last time step cannot accurately describe the success of different strategies in invading and establishing in the North patch. We therefore used the long-term temporal mean population size of each reproductive strategy as a measurement of their ecological success. Furthermore, we compared the long-term temporal mean with the initial short-term temporal mean during the first one hundred generations (corresponding to the first 10% of the total number of generations used in the simulations). This comparison between long-term and initial temporal mean population sizes can indicate whether a particular reproductive strategy is more successful at invading an unstable environment and/or outcompeting a competing strategy in the long term.

In the competition simulations, we assumed that the South patch hosts a large 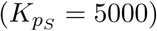 ancestral population composed of individuals that reproduce using different strategies, while the North patch is a smaller 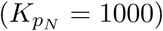, empty marginal habitat that is open and available for colonization. The North patch is partially connected to the South patch such that individuals can migrate between patches at a specified rate. Environmental conditions in the South patch are stable and do not change (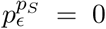 and 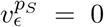), while the North patch experiences environmental stress at different levels. For all initial ancestral populations tested, we explored the effect of different environmental regimes in the North patch, in terms of probability of occurence of stressful environmental conditions 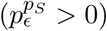 and maximum level of stress 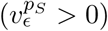.

Different compositions of the ancestral population were used in order to explore the interaction effects of different strategies during habitat invasion and colonization. We analyzed the pairwise dynamics of habitat colonization in the North patch by setting ancestral populations composed of two different strategies for each possible combination of strategies, allowing us to investigate the effect of each strategy on each other strategy when they compete for dominance during invasion of the North patch. In the initial conditions, competing reproductive strategies in the ancestral populations were equally represented in terms of initial number of individuals 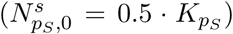. Because the initial subpopulation size of each strategy in the ancestral population is above the Allee threshold 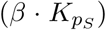, all strategies have the same initial growth rate in the South patch. All simulations were performed on R version 3.5.2 [47].

## Results

When the ancestral population in the South patch remains isolated (no migration between patches) and environmental conditions are stable (no environmental stress), population structure (in terms of proportion of the population composed by each reproductive strategy) remains constant and no particular reproductive strategy is ecologically more successful than any competing strategy in terms of temporal mean population size. When the effect of density is ignored, strategies that reproduce asexually (A, B, D- and E-) have their maximum growth rate because environmental conditions are optimal and strategies that reproduce sexually (C, D+ and E+) have their maximum growth rate both because environmental conditions are optimal and the subpopulation size is greater than the Allee saturation size. Therefore, sexual and asexual reproduction are equally successful under the baseline conditions present in the ancestral population.

In order to assess initial invasion and long-term ecological success of different reproductive strategies in the North patch, we calculated the temporal mean population size of the competing strategies during the first one hundred generations and in the long term (over the total simulation time). We then systematically analyzed the success of different strategies in different environments by changing the values of the parameters that control the Allee effect (*α* and *β*) as well as environmental stress (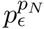 and 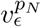) and sensitivity to environmental stress (*ϕ*^*B*^ and *ϕ*^*C*^).

### Population dynamics during invasion of a marginal, unstable habitat

When a marginal, unstable habitat (North patch) becomes available, migration from the ancestral population (South patch) to the North patch drives the initial invasion of the North patch by both competing strategies proportionally to their frequencies in the ancestral population. Nonetheless, because the Northern population faces environmental conditions that are different from the conditions in the South patch, the population structure in the North patch diverges considerably from that in the South patch (e.g., Figure S4). Despite fluctuations in population size and composition due to environmental stress events, the population composition in the North patch reaches a clear pattern in terms of frequency dominance of different reproductive strategies for different environmental conditions. Note that, in many cases, strategies that perform better during the initial stages of invasion are not always the most successful in the long term (explained in more detail below).

Furthermore, the population structure in the North patch can affect the population structure in the South patch through migration. However, because the population size in the North patch is much smaller than in the South patch 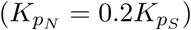 and migration from the North patch to the South patch occurs at a lower rate than in the opposite direction, changes in the population structure in the South patch occurs at a much lower rate (e.g., Figure S5). This lower rate of change can be explained by the small effect of the number of incoming individuals to the South patch relative to its population size.

### Allee effect and environmental stress

In a general case, we simulated pairwise competitions between different strategies, focusing on the Allee effect under two environmental scenarios where the probability of environmental stress is 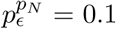 and stress levels can be either low (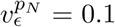; Figures S6-S7) or high (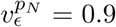 Figures S8-S9). We then simulated pairwise competitions focusing on environmental variation when sexuals are under weak (*α* = 0.3 and *β* = 0.1; Figures S10-S11) or strong (*α* = 0.1 and *β* = 0.3; Figures S12-S13) Allee effects.

In our model, *α* and *β* determine the Allee effect curvature and saturation point, respectively, and therefore can explain some of the differences in ecological success between parthenogens (strategies A, B, D- and E-) and sexuals (strategies C, D+ and E+). According to our Allee function, the Allee effect becomes stronger when *α* → 0^+^ and *β* → ∞; and weaker when *α* → 0^−^ and *β* → 0. This can be observed in Figures S6-S9. The Allee effect is particularly important during the initial invasion, when sexuals struggle to reproduce while parthenogens thrive under low environmental stress. However, parthenogens are particularly sensitive to the probability of encountering stressful conditions 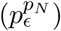 and the level of stress experienced 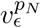, often allowing sexuals to outcompete them under stressful conditions even when the Allee effect is moderate (Figures S10-S13). In the long term, even when the Allee effect is strong, sexuals can outcompete parthenogens, showing a greater temporal mean population size. Because of the complex interactions between sensitivity to environmental stress and Allee effect, we analyzed each pairwise competition separately focusing on the biological properties of each strategy (below).

### Obligate apomictic vs. obligate automictic parthenogenesis

Strategy A (obligate apomictic parthenogenesis) was assumed to have maximum sensitivity to environmental stress (*ϕ*^*A*^ = 1.0), while strategies B (obligate automictic parthenogenesis) and C (obligate sexual reproduction) were assumed to have a lower sensitivity (*ϕ*^*A*^ > *ϕ*^*B*^ > *ϕ*^*C*^). Since apomictic and automictic parthenogens differ only in the magnitude of their response to environmental stress, competition between these strategies is expected to favor the dominance of automictic parthenogens (strategy B), and lower sensitivity (*ϕ*^*B*^ ≪ *ϕ*^*A*^) leads to a greater dominance of automictic parthenogens in the population (Figures S14-S15). This is a consequence of the assumption that automictic parthenogenesis has no extra fitness cost relative to apomictic parthenogenesis but generates phenotypic diversity which reduces the sensitivity of the population to environmental stress. This difference in temporal mean subpopulation size of apomictic and automictic parthenogens is particularly strong under highly stressful conditions 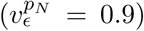. However, although our model assumes there is a sensitivity difference between apomictic and automictic parthenogenesis, empirical data quantifying the magnitude of this difference is absent. Additionally, fitness costs of automictic parthenogenesis relative to apomictic parthenogenesis remain unexplored.

### Obligate parthenogenesis vs. obligate sexual reproduction

As mentioned above, growth rate of obligate sexuals is affected not only by their sensitivity to environmental stress but also by the Allee effect caused by the difficulty in finding compatible mating partners when the population size is small. If the Allee effect is weak (*α* = 0.3 and *β* = 0.1) during the initial invasion of the marginal habitat, parthenogenetic reproduction is particularly favored when environmental stress is low 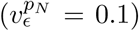, while sexual reproduction is particularly favored under high environmental stress 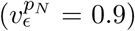 as long as their sensitivity is low enough to overcome the Allee effect (Figure S16). In the long term, sexuals outperform parthenogens even when their sensitivity to environmental stress (*ϕ*^*C*^ = 0.9) approaches that of parthenogens, although in such cases the difference in ecological success is small (Figure S17). If the Allee effect is strong (*α* = 0.1 and *β* = 0.3), however, parthenogens (*ϕ*^*A*^ = 1.0) are generally more successful than sexuals during the initial invasion and in the long term, except when sexuals are considerably less sensitive than parthenogens under very stressful environmental conditions 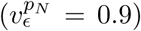, significantly reducing the population growth rate of parthenogenetic invaders (Figure 3-4).

**Figure 3:**
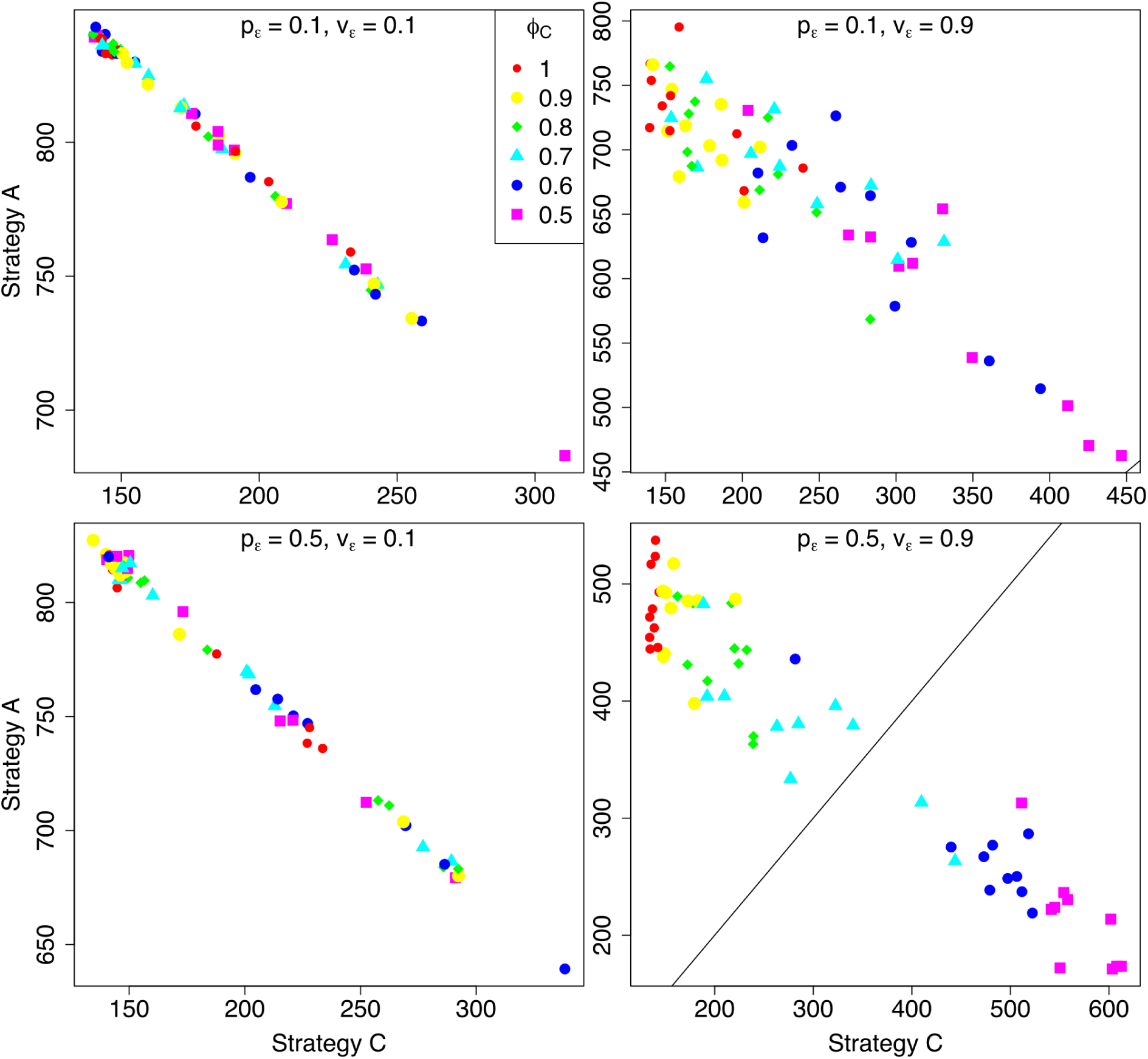
Obligate parthenogenesis (strategy A/B) vs. obligate sexual reproduction (strategy C). Axes represent the short-term temporal mean population sizes of competing strategies in the marginal habitat (North path) under a strong Allee effect (*α* = 0.1 and *β* = 0.3). Note that *ϕ* represents the relative difference in sensitivity to environmental stress between apomictic (strategy A; *ϕ*^*A*^ = 1.0) and automictic (strategy B; *ϕ*^*B*^) parthenogenesis, or between parthenogenesis (strategy A; *ϕ*^*A*^ = 1.0) and sexual reproduction (strategy C; *ϕ*^*C*^).

**Figure 4:**
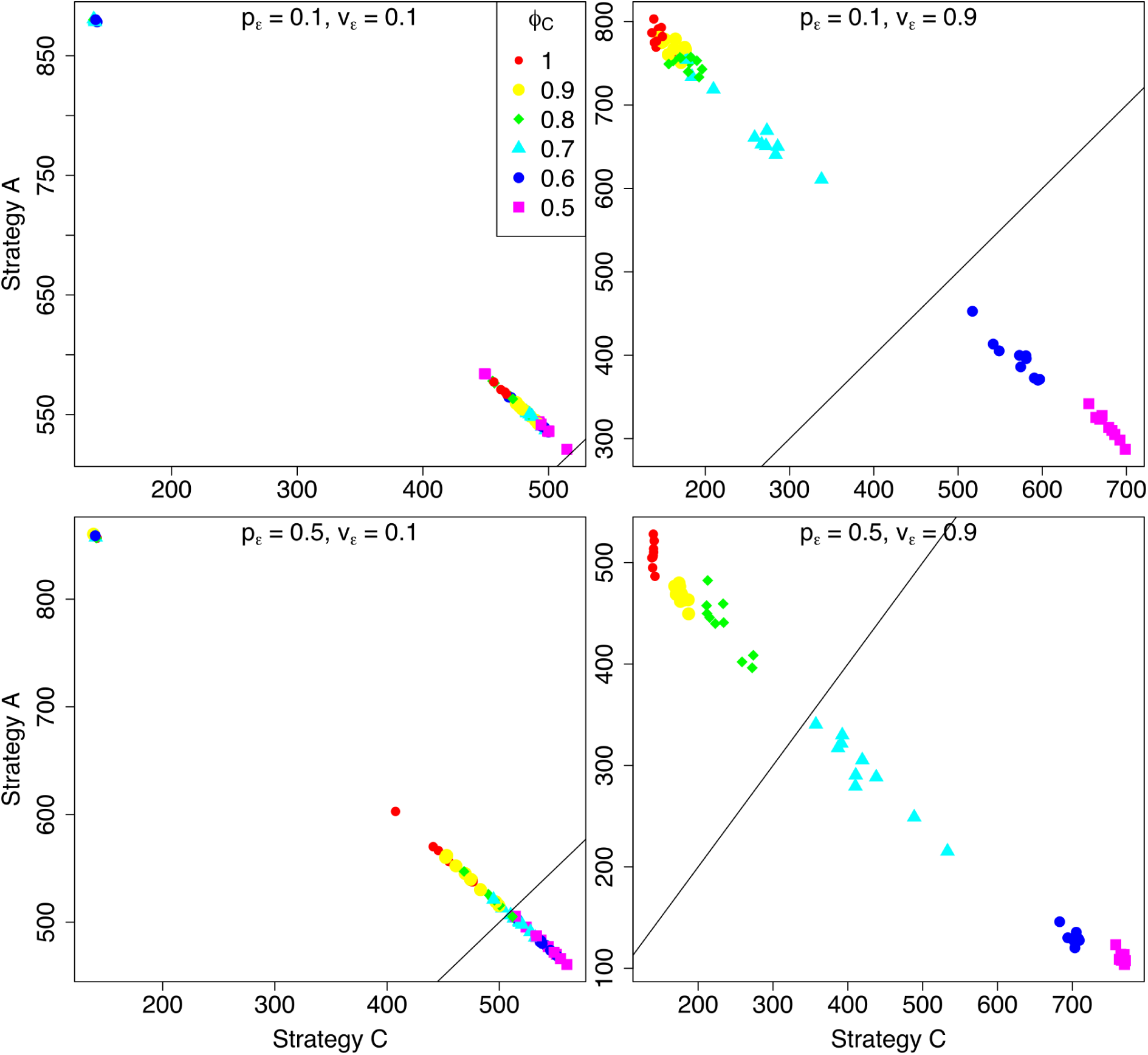
Obligate parthenogenesis (strategy A/B) vs. obligate sexual reproduction (strategy C). Axes represent the long-term temporal mean population sizes of competing strategies in the marginal habitat (North path) under a strong Allee effect (*α* = 0.1 and *β* = 0.3). Note that *ϕ* represents the relative difference in sensitivity to environmental stress between apomictic (strategy A; *ϕ*^*A*^ = 1.0) and automictic (strategy B; *ϕ*^*B*^) parthenogenesis, or between parthenogenesis (strategy A; *ϕ*^*A*^ = 1.0) and sexual reproduction (strategy C; *ϕ*^*C*^).

In a deterministic simulation, with a constant environmental stress level in the marginal habitat (Δ*ϵ* ∈ {0.1, 0.5, 0.9}) and constant migration rates (*ρ* = 0), increasing the magnitude of the Allee effect (*α* → 0^+^; *β* = 0.3) and the environmental sensitivity of sexuals (*ϕ*^*C*^) relative to parthenogens (*ϕ*^*C*^ approaches *ϕ*^*A*^) increases the temporal dominance of parthenogens in the marginal habitat (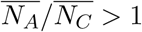; Figure 5). However, this effect is weaker in the long term.

**Figure 5:**
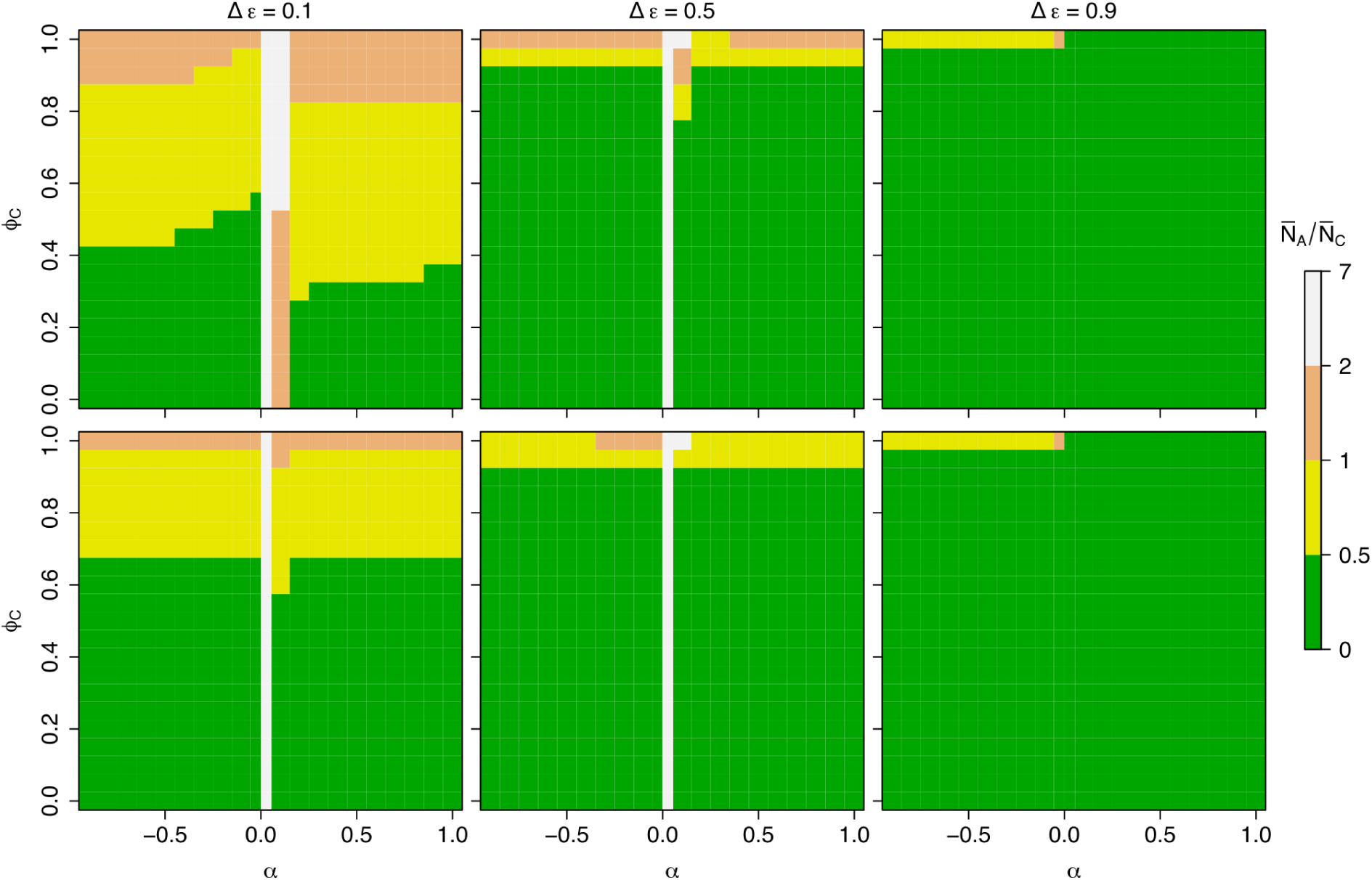
Obligate parthenogenesis (strategy A/B) vs. obligate sexual reproduction (strategy C). Deterministic ratio of short-term (top row) and long-term (bottom row) mean population sizes of obligate parthenogens to obligate sexuals 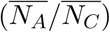 under constant environmental stress levels in the marginal habitat (Δ*ϵ* ∈ {0.1, 0.5, 0.9}), constant migration rates (*ρ* = 0), different values of *α* (Allee curve; *β* = 0.3) and different environmental sensitivity values of sexuals relative to parthenogens (variable *ϕ*^*C*^, with constant *ϕ*^*A*^ = 1.0).

In general, sexuals become less successful than parthenogens when *β* is large (e.g., *β* = 0.3), demanding a greater population size in order to reach Allee saturation 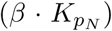, and *α* is positive and close to zero (*α* = 0.1, in our simulations), demanding a greater population size in order to reach the Allee threshold (Equation 3). If the Allee threshold and saturation point are large, sexuals will only be more successful than parthenogens under three conditions: over a very large number of generations (assuming no extinction; > 1000 generations); when environmental stress levels are high such that parthenogens are affected more strongly than sexuals under a strong Allee effect; or when the *per capita* growth rate is high enough for the Allee threshold and saturation point to be reached in a relatively short period of time (not shown here). These results show that, for obligate sexuals (strategy C) to outcompete obligate parthenogens (strategies A and B), there must be a balance between their sensitivity to environmental stress and the Allee effect that they are subject to such that the net growth rate of obligate sexuals becomes greater than the net growth rate of obligate parthenogens.

### Facultative parthenogenesis vs. obligate sexual reproduction

Among sexually reproducing strategies, facultative parthenogens (strategy D/E) can perform significantly better than obligate sexuals (strategy C) because of their ability to reproduce asexually when the population size is small, partially avoiding the Allee effect. Under a weak Allee effect, both strategies are equally successful (Figures S18-S19). Differences are visibly significant when the Allee effect is strong (Figures 6-7) and transitions from sexual to parthenogenetic reproduction become particularly advantageous during the initial invasion. Furthermore, the difference in temporal mean population size between strategies increases as sexuals become more sensitive to environmental stress, reducing the advantage of sex even further especially under stressful conditions. A higher transition rate *τ* can also increase the success of facultative parthenogens, but the increase is only visible under stressful environmental conditions and strong Allee effect on sexuals (Figures S20-S21).

**Figure 6:**
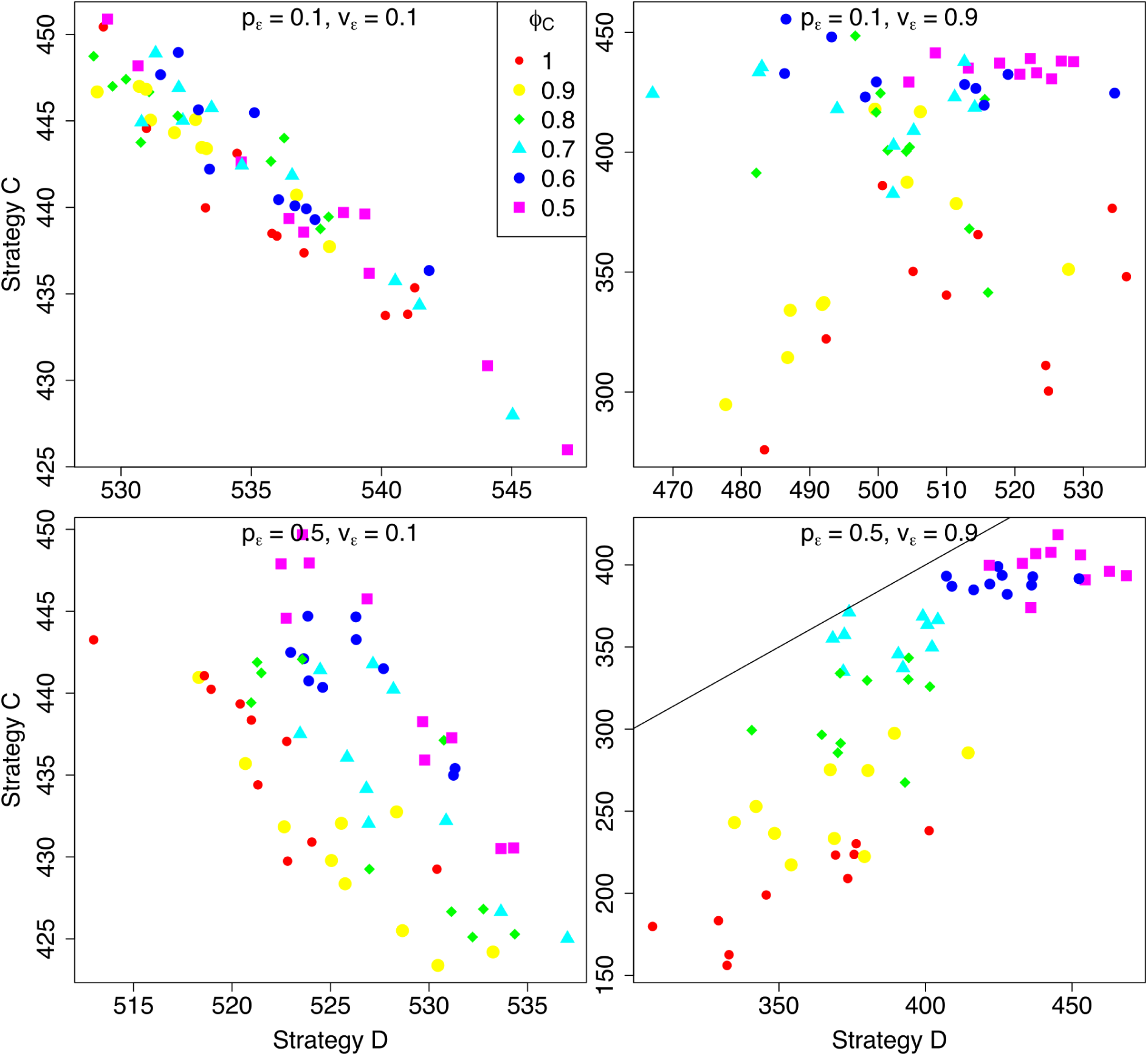
Obligate sexual reproduction (strategy C) vs. facultative parthenogenesis (strategy D/E). Axes represent the short-term temporal mean population sizes of competing strategies in the marginal habitat (North path) under a strong Allee effect (*α* = 0.1 and *β* = 0.3). Note that *ϕ* represents the relative difference in sensitivity to environmental stress between apomictic (strategy A; *ϕ*^*A*^ = 1.0) and automictic (strategy B; *ϕ*^*B*^) parthenogenesis, or between parthenogenesis (strategy A; *ϕ*^*A*^ = 1.0) and sexual reproduction (strategy C; *ϕ*^*C*^).

**Figure 7:**
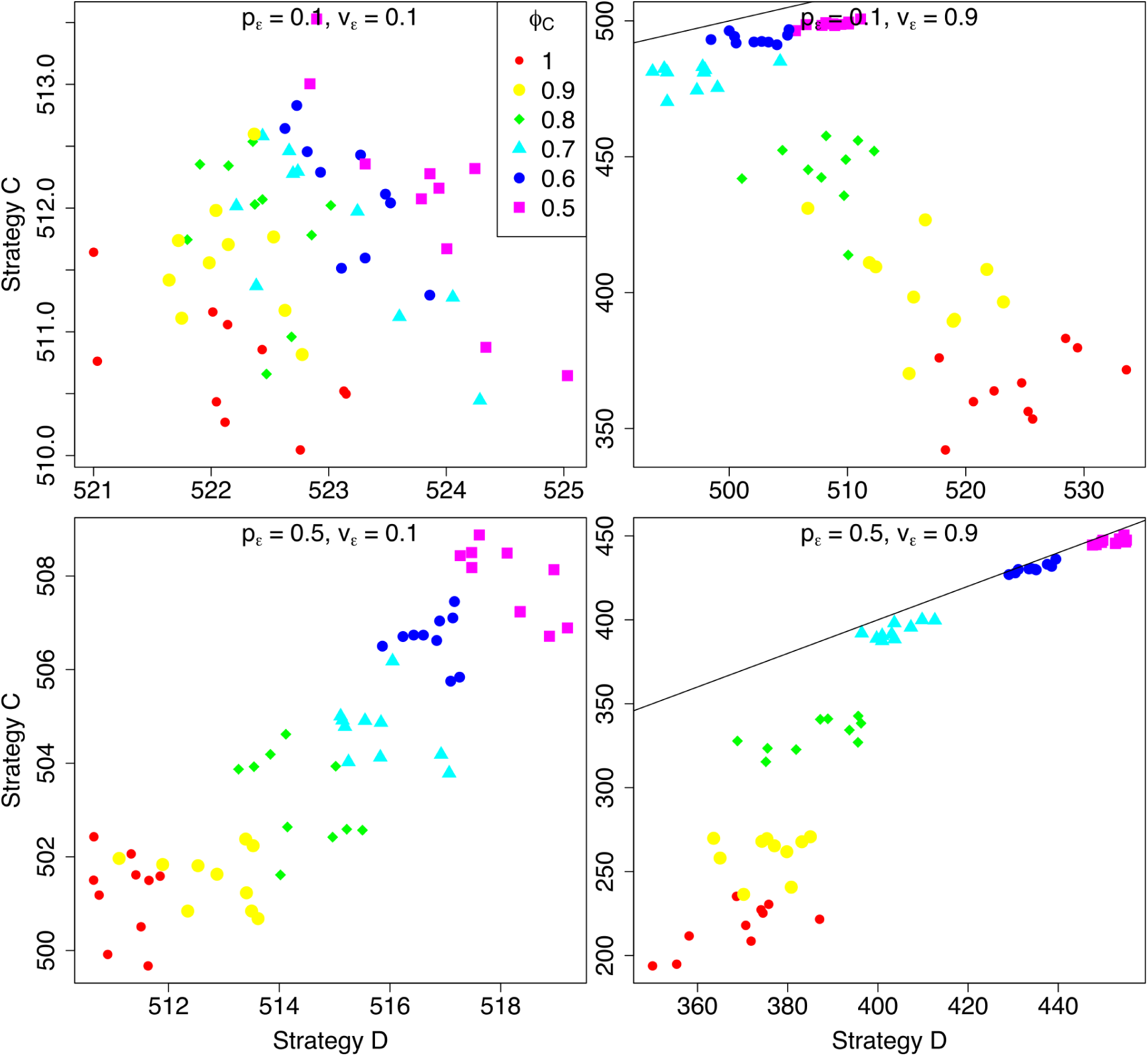
Obligate sexual reproduction (strategy C) vs. facultative parthenogenesis (strategy D/E). Axes represent the long-term temporal mean population sizes of competing strategies in the marginal habitat (North path) under a strong Allee effect (*α* = 0.1 and *β* = 0.3). Note that *ϕ* represents the relative difference in sensitivity to environmental stress between apomictic (strategy A; *ϕ*^*A*^ = 1.0) and automictic (strategy B; *ϕ*^*B*^) parthenogenesis, or between parthenogenesis (strategy A; *ϕ*^*A*^ = 1.0) and sexual reproduction (strategy C; *ϕ*^*C*^).

### Obligate parthenogenesis vs. facultative parthenogenesis

As mentioned above, facultative parthenogens (strategy D/E) reproduce sexually in the ancestral population because of the large population size and high environmental stability, so incoming migrants to the marginal habitat are subject to the Allee effect during the very first wave of migration, which can explain their lower ecological success relative to obligate parthenogens (strategy A/B) when sexuals and parthenogens have similar sensitivities to environmnetal stress (Figures S22-S25). When sexuals are considerably less sensitive, the advantage of sex can compensate for the Allee effect and lead to a higher ecological success of facultative parthenogens even under a strong Allee effect. The balance between sensitivity to environmental stress and the Allee effect that equalizes the success of sexuals (including facultative parthenogens) and obligate parthenogens depends on the environmental conditions experiences. For example, under highly stressful environmental conditions (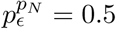 and 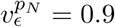), sexuals under a strong Allee effect must be relatively less sensitive than sexuals under a weak Allee effect in order to outcompete parthenogens.

## Discussion

Our results show that the relative frequency of a reproductive strategy in a marginal habitat can be determined by a complex interaction between the environmental conditions in that habitat, the magnitude of the Allee effect and the relative sensitivity of competing strategies to environmental stress. These factors may explain the empirically observed biased distribution of sexuals and asexuals (parthenogens) along a habitat range, with asexuals being particularly abundant in marginal habitats, a pattern that characterizes geographic parthenogenesis. In our model, an equal maximum intrinsic growth rate was assumed for sexual and asexual reproduction and, therefore, our results shed light on the importance of ecological (non-reproductive) processes as explanatory causes of geographic parthenogenesis.

### Geographic parthenogenesis and range expansion

Previous models explored different processes that can lead to geographic parthenogenesis. A spatially explicit genetic model showed that asexuals may be favored by local adaptation to a marginal habitat through clone selection and protection of optimal genotypes from gene flow from the core habitat [20]. Geographic parthenogenesis can also result from selection for resistance and high fecundity in a facultatively parthenogenetic metapopulation occupying an environmental gradient, limiting mating at the edge of the habitat and forcing females to reproduce asexually, generating a female bias [40]. However, the ecological success of asexual reproduction certainly depends on the environmental conditions and the characteristics of the competing reproductive strategy. A model exploring the frozen niche variation hypothesis [48] indicated that, despite its two-fold advantage relative to sexual reproduction [2], asexual clonal reproduction may have its invasion probability reduced due to a fast accumulation of deleterious mutations in the initially small clonal population [49], decreasing its initial advantage of local adaptation. Clonal invasion probability can be reduced even further when sexual populations have many niche phenotypes, requiring the occurence of a beneficial mutation for niche exploration within the asexual clonal population before individuals can invade a niche occupied by sexuals [49]. Additionally, it has been proposed that metapopulation dynamics in marginal habitats may be detrimental to sexuals because they may suffer more intensely from genetic bottlenecks and inbreeding [16].

Although most studies focus on the coexistence or elimination of reproductive strategies as a result of active selection leading to evolutionary change, it has been shown theoretically that stochastic demographic processes may lead to geographic structure in the distribution of sexual and asexual morphs in recently invaded areas without having to invoke adaptive differences. However, these patterns (e.g. clone dominance) are transient and are eventually substituted by sexual reproduction [27]. Other models explored the distribution of individuals within a habitat and the factors that determine population range limits, although not in the context of geographic parthenogenesis. However, because parthenogens are typically found in marginal habitats, the study of range limits can be particularly useful for understanding the geography of reproductive strategies. Range limits may have many causes, each leading to different evolutionary outcomes [50], with range expansion being typically initiated through dispersal and subsequent niche evolution in the new habitat. This process of dispersal leading to range expansion has been explored in the context of a species invasion of a novel habitat, where niche evolution has been found to be affected by several factors, including the initial maladaptation of the invading population, mutation rate and degree of heterogeneity in the occupied range [51]. These factors, as mentioned in the present study, can affect different reproductive strategies differently and therefore lead to the observed biased geographic distribution at the population range limits (e.g., [52]). The Allee effect on sexual populations, for example, can explain the difference in the spatial distribution of sexuals and asexuals during range expansion.

### The Allee effect on population dynamics

The Allee effect explored in our model reduces the growth rate of sexual populations when the population size is small. This effect has been observed in many populations and has been suggested to cause species extinctions [53; 54]. For example, experimental observations from the annual herb *Clarkia concinna* suggest that small populations are more likely to go extinct because of the Allee effect caused by the lack of effective pollination, leading to reproductive failure [55]. In populations of the shrub *Banksia goodii*, there is a clear positive relationship between population size, number of seeds per unit population size and fraction of fertile plants, with very small populations producing a disproportionately small number of seeds, a pattern that resembles the strong Allee effect (*α* = 0.1 and *β* = 0.3) used in our model and that can lead to local extinction [56]. In natural populations of the Glanville fritillary butterfly *Melitaea cinxia*, the fraction of mated females decreases with decreasing local population density, resulting in a reduced reproductive success in small populations [57]. This density-dependent growth rate has also been detected in many populations of the Atlantic herring *Clupea harengus* [58] and even in the bacterium *Vibrio fischeri* [59]. In the latter study, experiments using different initial population sizes of *Vibrio fischeri* showed a non-linear positive relationship between initial population density and the proportion of populations establishing in the media at the time of measurement, indicating a reduction in population growth rate when population density is low [59].

This positive density-dependent reproductive rate has also been explored from a theoretical perspective. The Allee effect was used in a model to predict the rate of population spread in the house finch *Carpodacus mexicanus* and density of birds near the center of the range[60], with a weak Allee effect, measured in terms of the fraction of birds mated as a function of population density, resembling the Allee curve with *α* = −0.1 in our model. A different model explored the Allee effect from the perpective of mate location dynamics, showing that very low population densities decrease the recognition of potential conspecific mates and therefore the probability of mating, driving the population to extinction [54]. The identification of conspecific mates is one of the factors that affects the Allee effect and should be taken into consideration in the future when calculating empirical values of *α* and *β* for the Allee curve given by Equation 2 in the present model.

More general models showed that the Allee effect may reduce the rate at which an invader moves to a new enviroment [61; 62] and the interaction between interspecific competition and the Allee effect can result in stability patterns that differ from models that ignore Allee effects [63; 64], which is also important for the competitive dynamics between different reproductive strategies explored in the current study. In the context of metapopulations, it has been suggested that the Allee effect may prevent small metapopulations from increasing even when resources are abundant or make large metapopulations go extinct due to stochastic environmental stress events when the number of occupied patches is small [65; 66]. Because of that, the effect of stressful events on population growth can affect the invasion of marginal habitats and subsequent range expansion.

### Environmental effects on population growth

Our model also considered the sensitivity of the population to environmental stress, which may depend on genetic and phenotypic diversity. Empirical studies in both animals and plants have shown that the founder genotypic and heritable phenotypic diversity is key to successful invasion, range expansion and establishment in new habitats where the population may find novel (stressful) environmental conditions [67]. An experimental study in the flour beetle *Tribolium castaneum* showed that the probability of a founding population going extict and the mean population size after several generations are, respectively, inversely proportional and directly proportional to the founding level of genetic variation [68]. In the clonal plant *Ranunculus reptans*, when introduced to previously unoccupied habitats and exposed to severe stressful conditions (flood and drought), populations founded by different genetic sources increased in abundance relative to populations founded by genetic monocultures [69]. Interestingly, studies analyzing genetic diversity as a function of distance to the core habitat (central population) show that central populations have significantly higher diversity than populations located at marginal habitats [70], indicating that marginal populations may be more sensitive to environmental stress and that genetic diversity may be restricted in such habitats.

It has been suggested that in populations where genetic diversity is very limited (e.g., clonal populations) adaptation to stressful environmental conditions can be difficult when relying on new mutations, leading populations to decline or go extinct [71]. This negative relationship between genetic diversity and risk of extinction affects parthenogenetic populations more strongly than sexual populations but this risk can be reduced when the parthenogenetic population is multiclonal instead of monoclonal [48]. In our model, parthenogenetic populations are more sensitive to environmental stress and thus more likely to decline under stressful conditions. However, because of the constant migration from the ancestral population to the marginal habitat, the dynamics in our model does not lead to local extinction. It is important to note that populations declining due to sensitivity to stressful environmental conditions can resume growing when new genetic variation is introduced from different sources, a process called evolutionary rescue [72; 73], which may be achieved via, for example, sexual reproduction in clonal populations of facultative parthenogens. Because increased asexual reproduction is common towards marginal habitats, those populations are also more likely to decline when challenged by changes in environmental conditions, which can make facultative parthenogenesis particularly beneficial. In the Baltic Sea, asexual recruitment seems to be very common in many macrophytic populations, but sexual recruitment is not completely absent [74], supporting the idea that the ability to transition between sexual and asexual reproduction is beneficial. Because of all these processes related to the effect of environmental stress on population growth, it is important that future studies provide empirical measurements of the correlation between genetic/phenotypic diversity and population sensitivity to different types of environmental stress.

## Conclusion

We used a quantitative approach to explore the ecological processes that can lead to geographic parthenogenesis and the invasion of new habitats by different reproductive strategies. We analyzed the Allee effect on sexual populations and the population sensitivity to environmental stress during the invasion of a marginal, unstable habitat to demonstrate that a complex interaction between the Allee effect, sensitivity to environmental stress and the environmental conditions can determine the relative success of competing reproductive strategies during the initial invasion and long-term establishment in the marginal habitat. In particular, sexuals need to compensate for the reduction in growth rate due to the Allee effect through a reduced sensitivity to environmental stress. However, the reduction in sensitivity is only strongly beneficial under highly stressful environmental conditions. Unfortunately, despite the empirical evidence for the Allee effect and differential sensitivity to environmental stress, empirical quantification of such processes remain scarce. Controlled and accurate quantification of the Allee effect on population growth in nature are difficult to obtain because many factors (both ecological and genetic) may affect the magnitude of the effect. We suggest that the following processes may play particularly important roles: (i) the distribution pattern of the immigrants (e.g., uniform vs. aggregated distribution) in the marginal habitat; (ii) migration rate, which can affect the speed at which the Allee threshold is reached; (iii) sexual selection (in particular, female choice, which can limit mating); and (iv) sociality, which can create an Allee effect even in asexual populations. Similarly, sensitivity to environmental stress is difficult to quantify because it is highly dependent on the type of stress and the genetic diversity of the population. The following factors may be particularly important for the quantification of sensitivity: (i) mutation rate (particularly important for clonal populations); (i) phenotypic diversity (e.g., apomictic vs. automictic parthenogenesis); and (iii) ability to respond (adaptively) to environmental conditions (phenotypic plasticity). All these processes can potentially explain the distribution patterns of different reproductive strategies and the present study suggests new patterns for empirical investigation.

## Acknowledgements

The authors would like to thank Kerstin Johannesson and the anonymous reviewers for their useful comments on an earlier version of this manuscript. WTAFS, PRJ and KCH were funded by BaltHealth, which has received funding from BONUS (Art. 185), funded jointly by the EU, Innovation Fund Denmark (grants 6180-00001B and 6180-00002B), Forschungszentrum Jülich GmbH, German Federal Ministry of Education and Research (grant FKZ 03F0767A), Academy of Finland (grant 311966) and Swedish Foundation for Strategic Environmental Research (MISTRA). KH was funded by CeMEB (The Linnaeus Centre for Marine Evolutionary Biology) at the University of Gothenburg, Sweden.

## Author contributions

WTAFS conceived the initial numerical model, wrote the script and ran the simulations. All authors contributed to the design and implementation of the research, the discussion of the results and the writing of the manuscript.

## Supplementary Information

**Figure S1:**
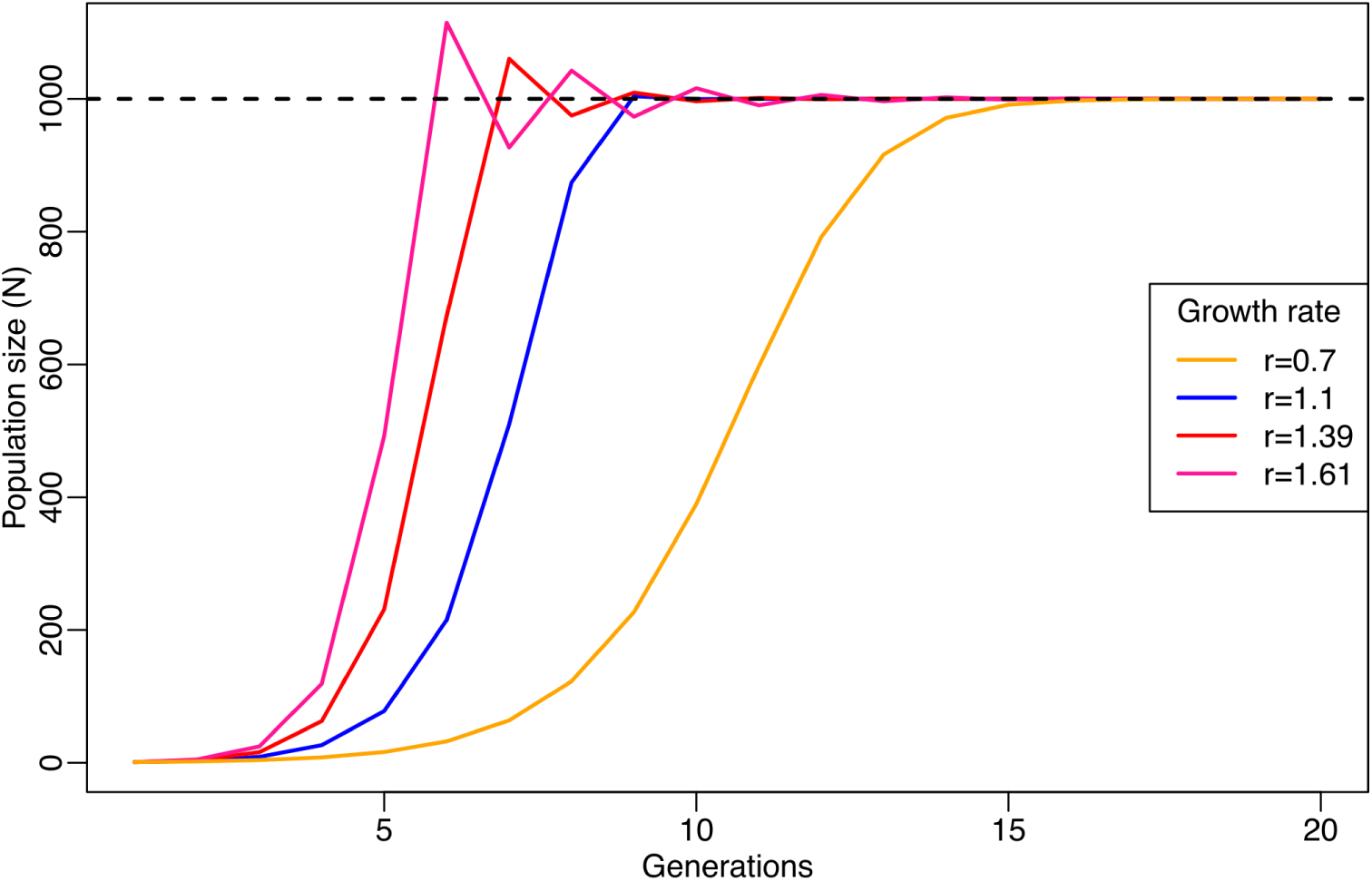
Logistic growth for different population growth rates. The maximum growth rate was set to *r*_*max*_ = 1.1 in all simulations.

**Figure S2:**
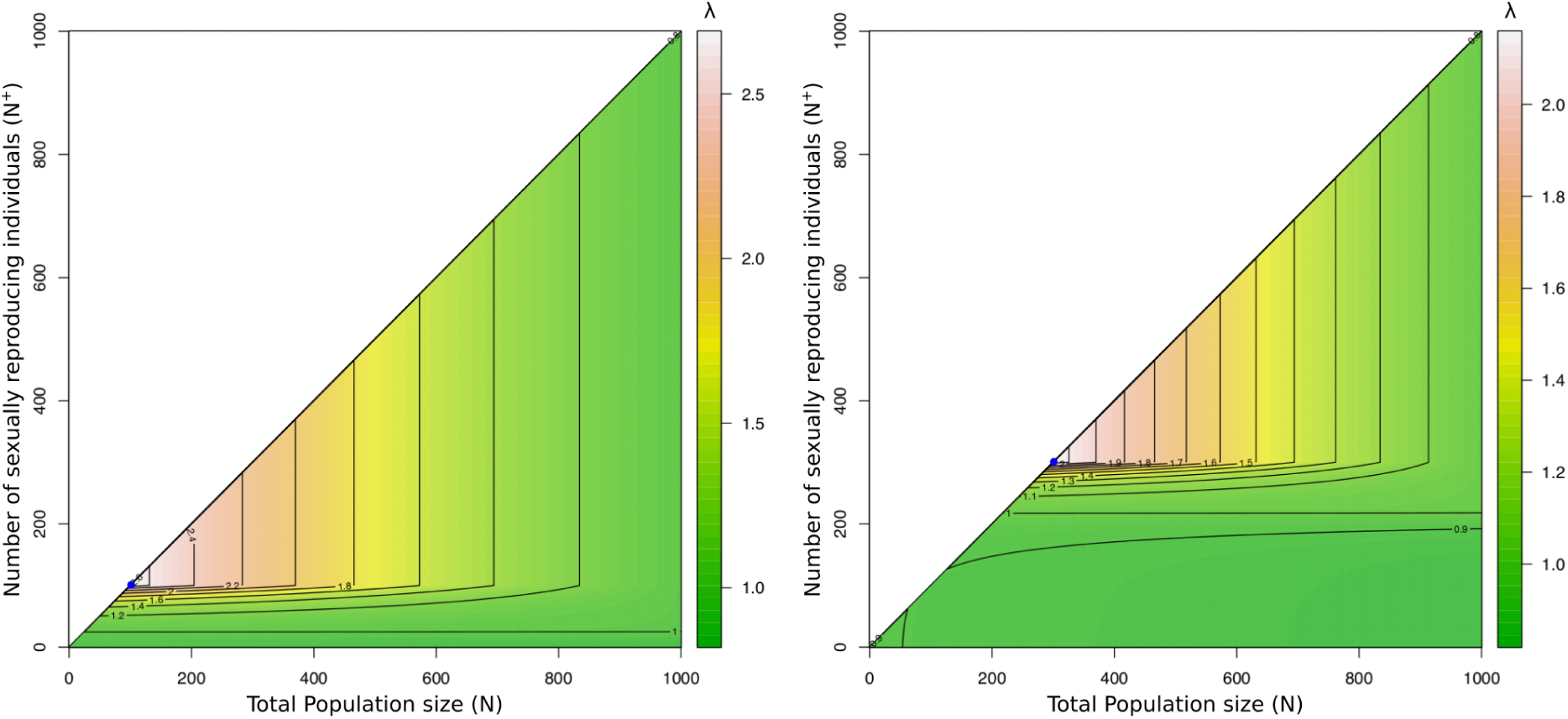
Density-dependent *per capita* growth rate (*λ*) of the sexually reproducing population under weak (*α* = 0.3 and *β* = 0.1; left) and strong (*α* = 0.1 and *β* = 0.3; right) Allee effects. Parameter values used across all simulations: *r*_*min*_ = −0.1, *r*_*max*_ = 1.1 and 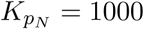.

**Figure S3:**
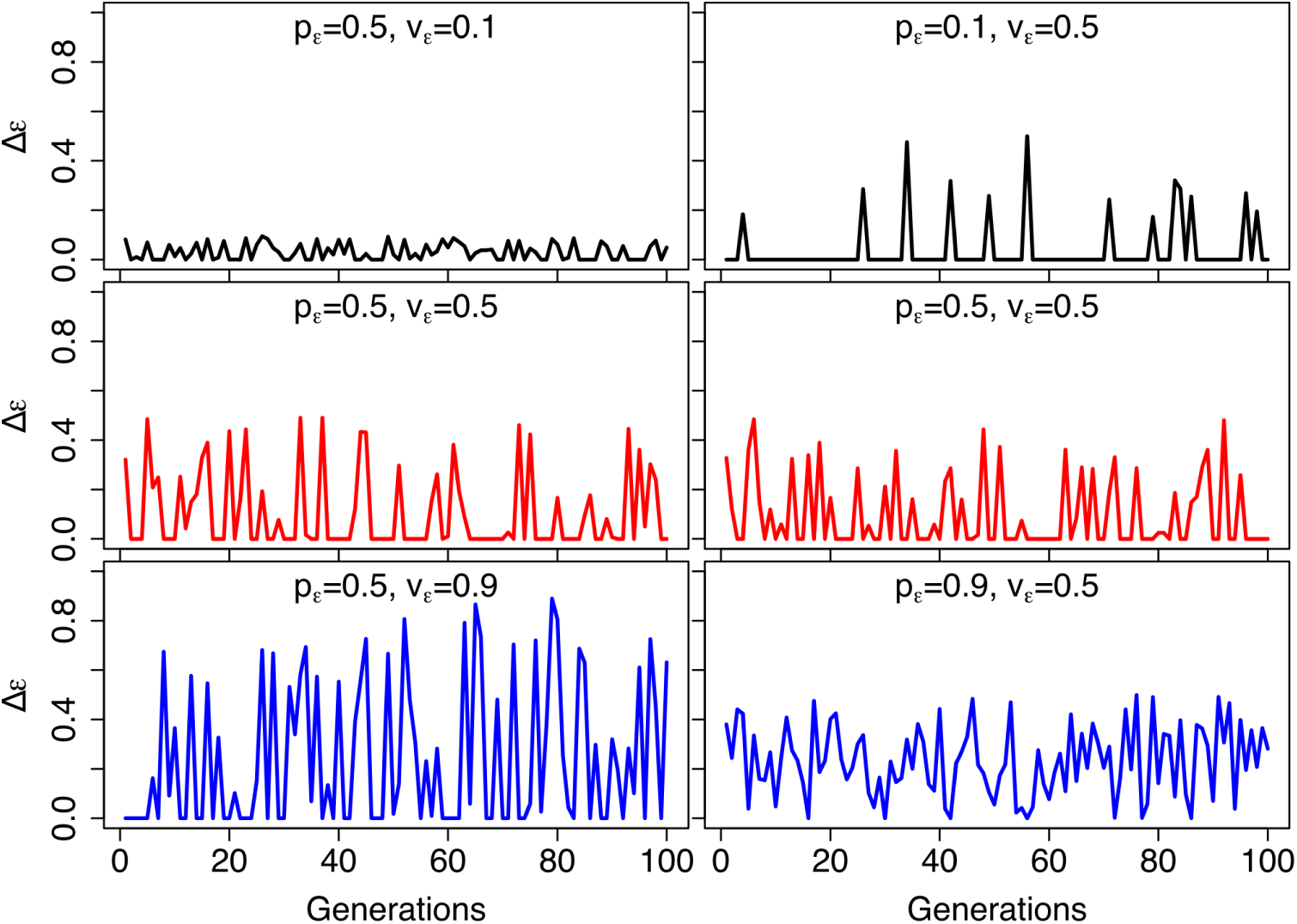
Examples of environmental dynamics showing environmental stress levels (Δ*ϵ*) for different probabilities of stress occurrence (*p*_*ϵ*_) and maximum stress level (*v*_*ϵ*_) throughout time.

**Figure S4:**
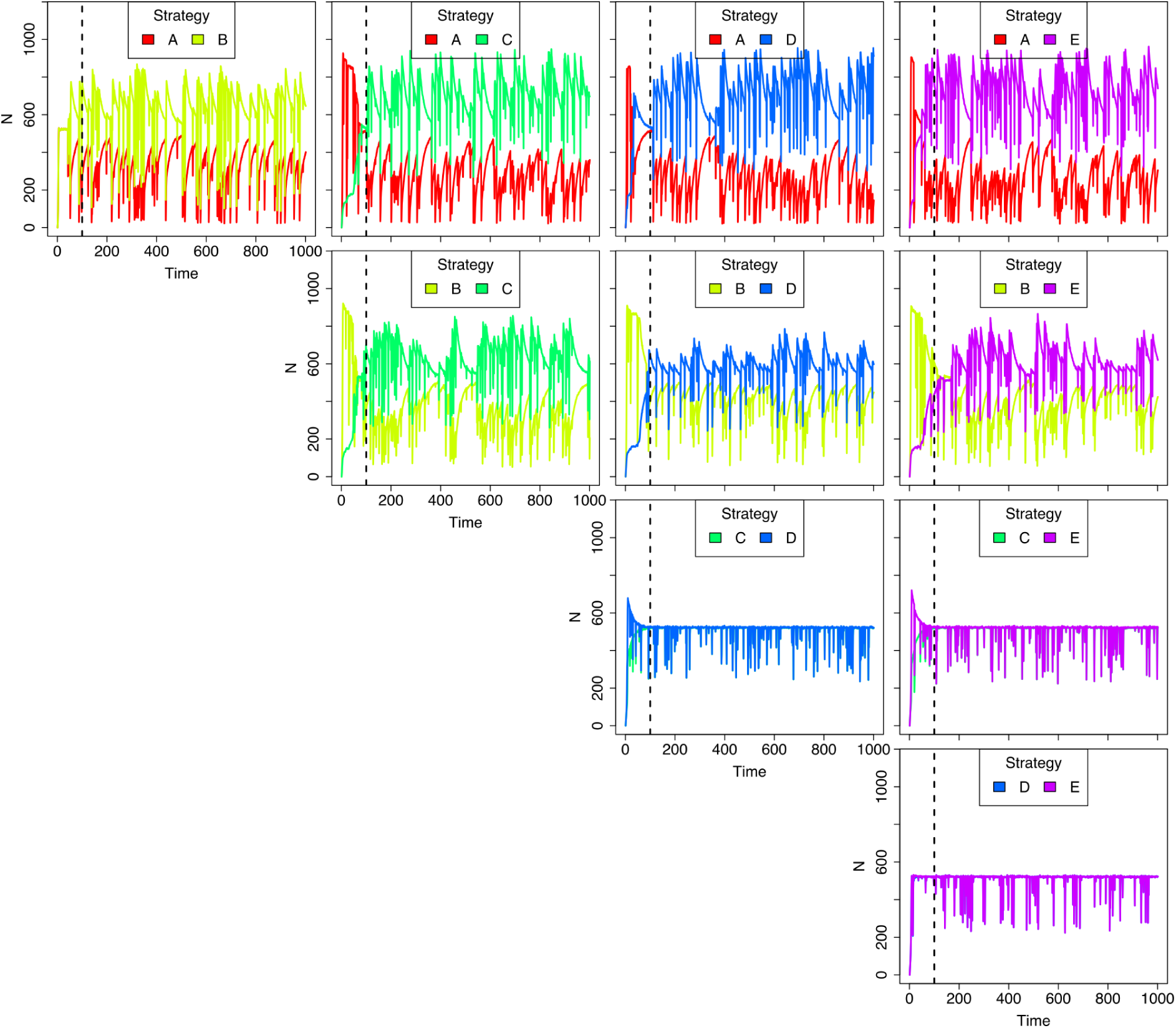
Pairwise competition between reproductive strategies and population dynamics during invasion of a marginal habitat (North patch) under high environmental stress (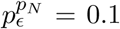 and 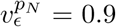), with sexual strategies exposed to a relatively strong Allee effect (*α* = 0.1 and *β* = 0.3). Dashed lines indicate generation 100. Sensitivity values: *ϕ*^*A*^ = 1.0, *ϕ*^*B*^ = 0.75, *ϕ*^*C*^ = 0.5.

**Figure S5:**
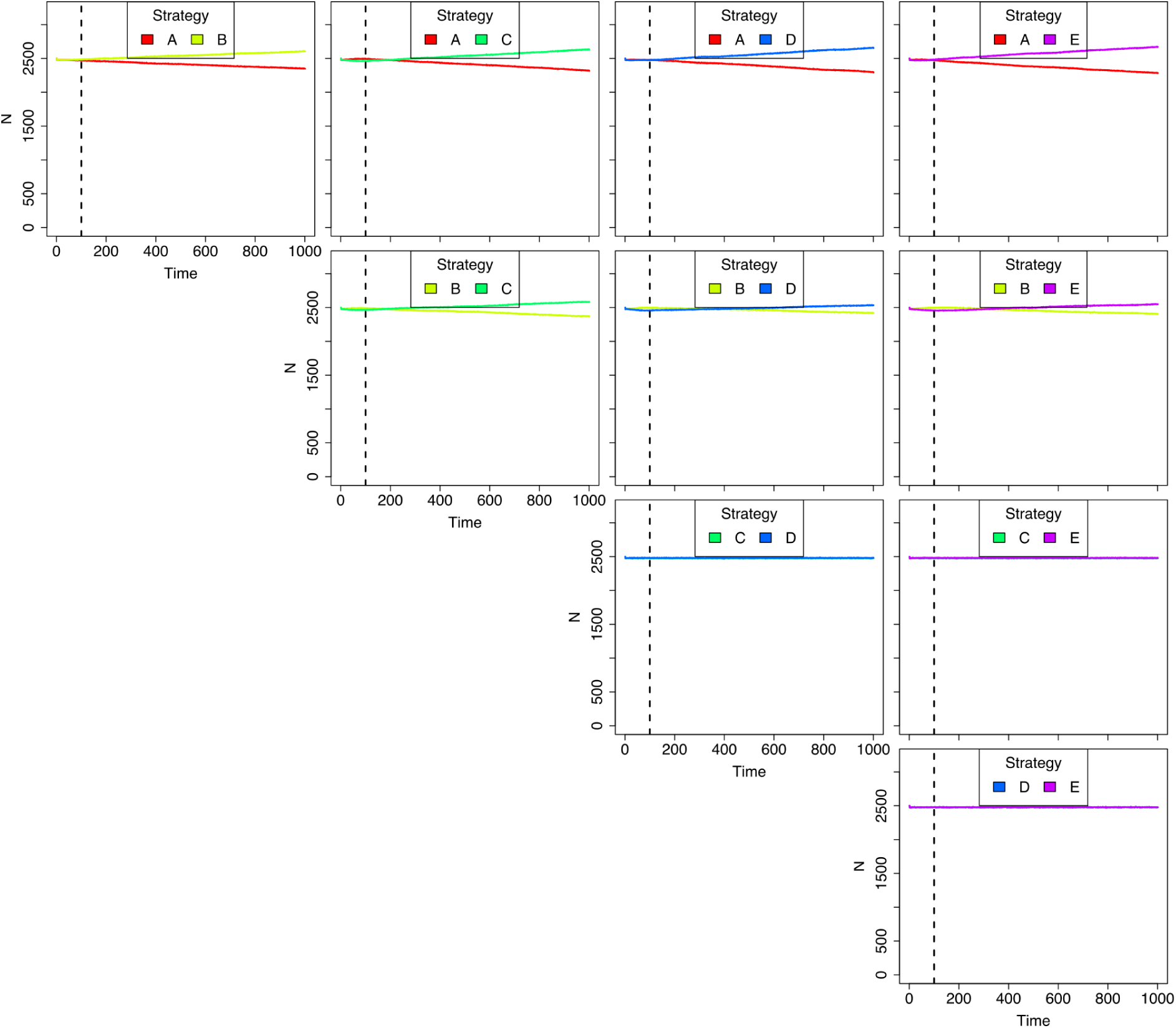
Pairwise competition between reproductive strategies and population dynamics in the ancestral population (South patch) during invasion of a marginal habitat (North patch) under high environmental stress (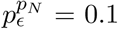 and 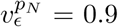), with sexual strategies exposed to a relatively strong Allee effect (*α* = 0.1 and *β* = 0.3). Dashed lines indicate generation 100. Sensitivity values: *ϕ*^*A*^ = 1.0, *ϕ*^*B*^ = 0.75, *ϕ*^*C*^ = 0.5.

**Figure S6:**
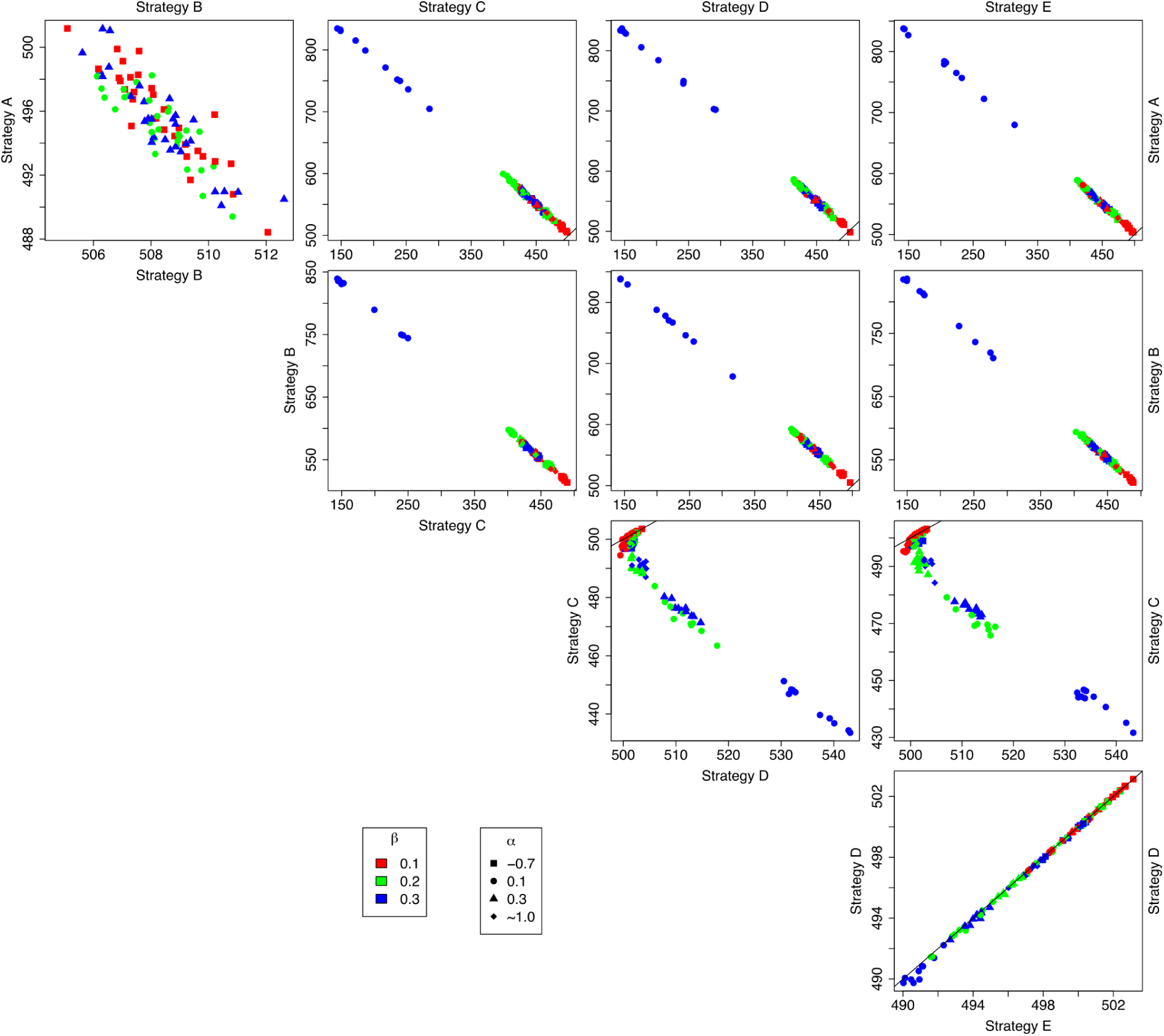
Pairwise competition between different reproductive strategies during invasion of a marginal habitat (North patch) under low environmental stress (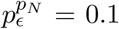 and 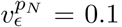), with sexual strategies exposed to a variety of Allee effect magnitudes (*α* and *β*). Axes represent the short-term temporal mean population sizes of competing strategies. Diagonal lines indicate identical temporal mean population sizes of competing reproductive strategies and therefore equivalent ecological successes. Sensitivity values: *ϕ*^*A*^ = 1.0, *ϕ*^*B*^ = 0.75, *ϕ*^*C*^ = 0.5.

**Figure S7:**
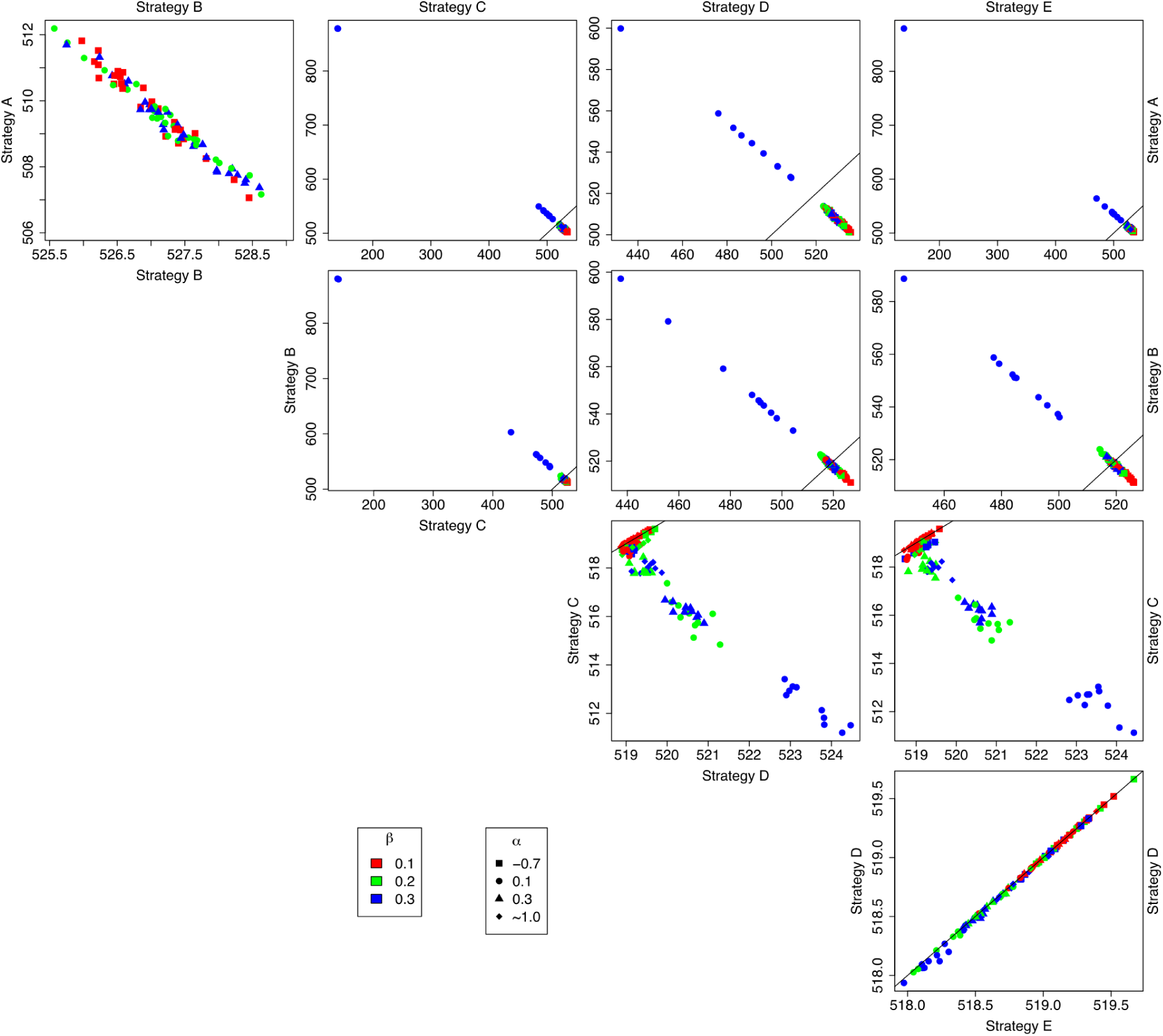
Pairwise competition between different reproductive strategies during invasion of a marginal habitat (North patch) under low environmental stress (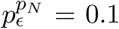 and 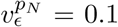), with sexual strategies exposed to a variety of Allee effect magnitudes (*α* and *β*). Axes represent the long-term temporal mean population sizes of competing strategies. Diagonal lines indicate identical temporal mean population sizes of competing reproductive strategies and therefore equivalent ecological successes. Sensitivity values: *ϕ*^*A*^ = 1.0, *ϕ*^*B*^ = 0.75, *ϕ*^*C*^ = 0.5.

**Figure S8:**
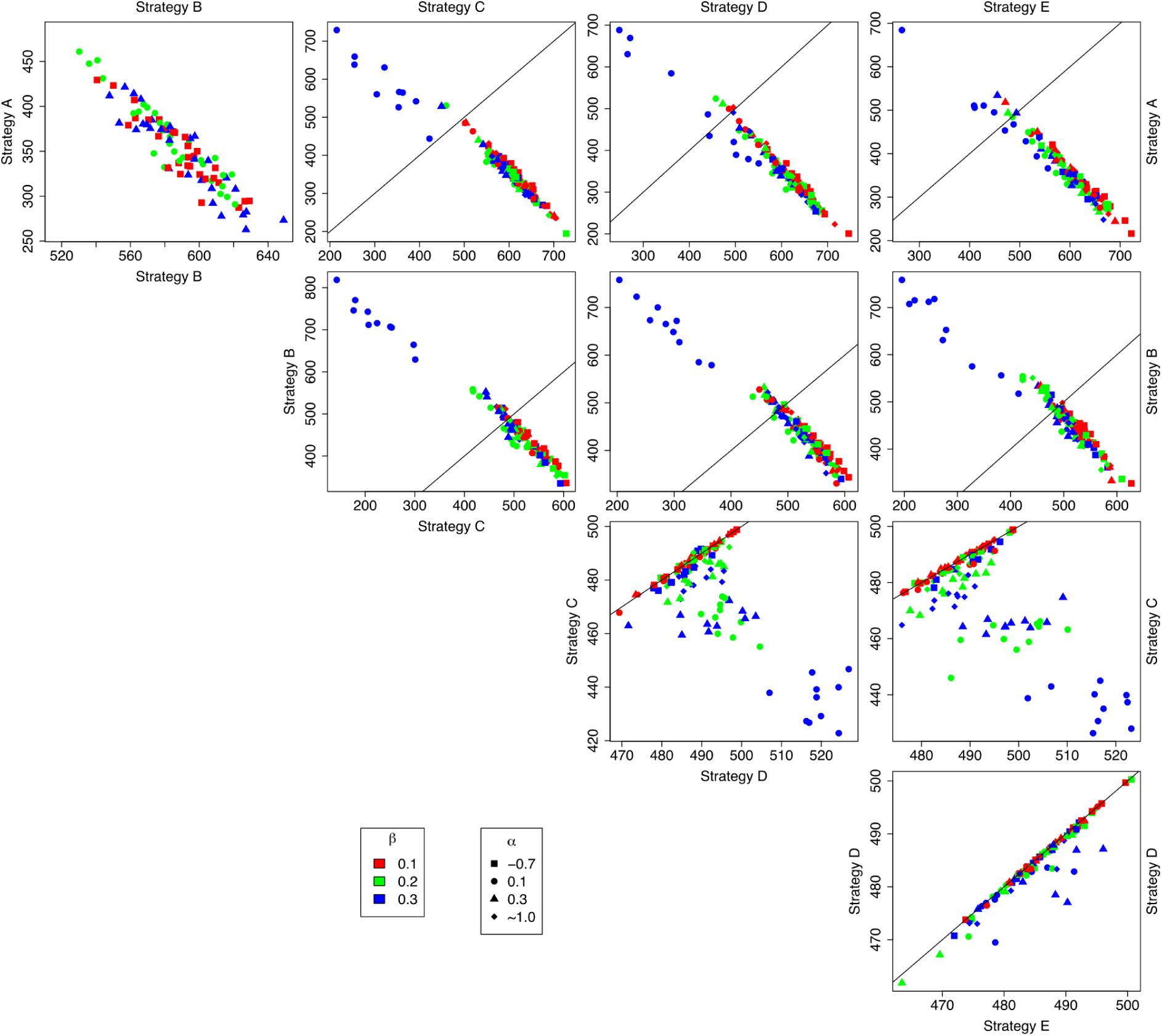
Pairwise competition between different reproductive strategies during invasion of a marginal habitat (North patch) under high environmental stress (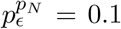 and 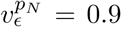), with sexual strategies exposed to a variety of Allee effect magnitudes (*α* and *β*). Axes represent the short-term temporal mean population sizes of competing strategies. Diagonal lines indicate identical temporal mean population sizes of competing reproductive strategies and therefore equivalent ecological successes. Sensitivity values: *ϕ*^*A*^ = 1.0, *ϕ*^*B*^ = 0.75, *ϕ*^*C*^ = 0.5.

**Figure S9:**
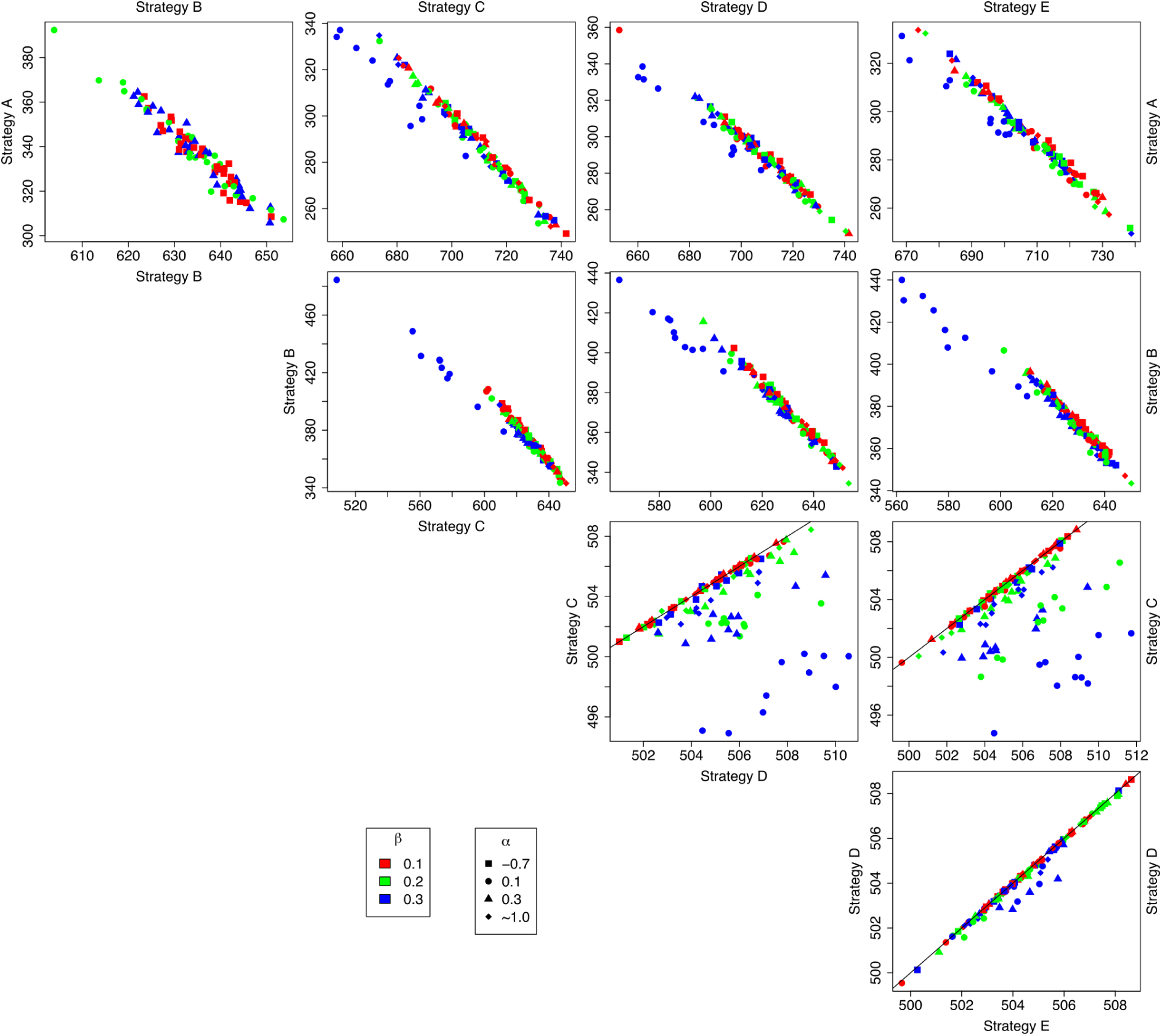
Pairwise competition between different reproductive strategies during invasion of a marginal habitat (North patch) under high environmental stress (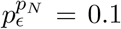 and 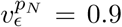),with sexual strategies exposed to a variety of Allee effect magnitudes (*α* and *β*). Axes represent the long-term temporal mean population sizes of competing strategies. Diagonal lines indicate identical temporal mean population sizes of competing reproductive strategies and therefore equivalent ecological successes. Sensitivity values: *ϕ*^*A*^ = 1.0, *ϕ*^*B*^ = 0.75, *ϕ*^*C*^ = 0.5.

**Figure S10:**
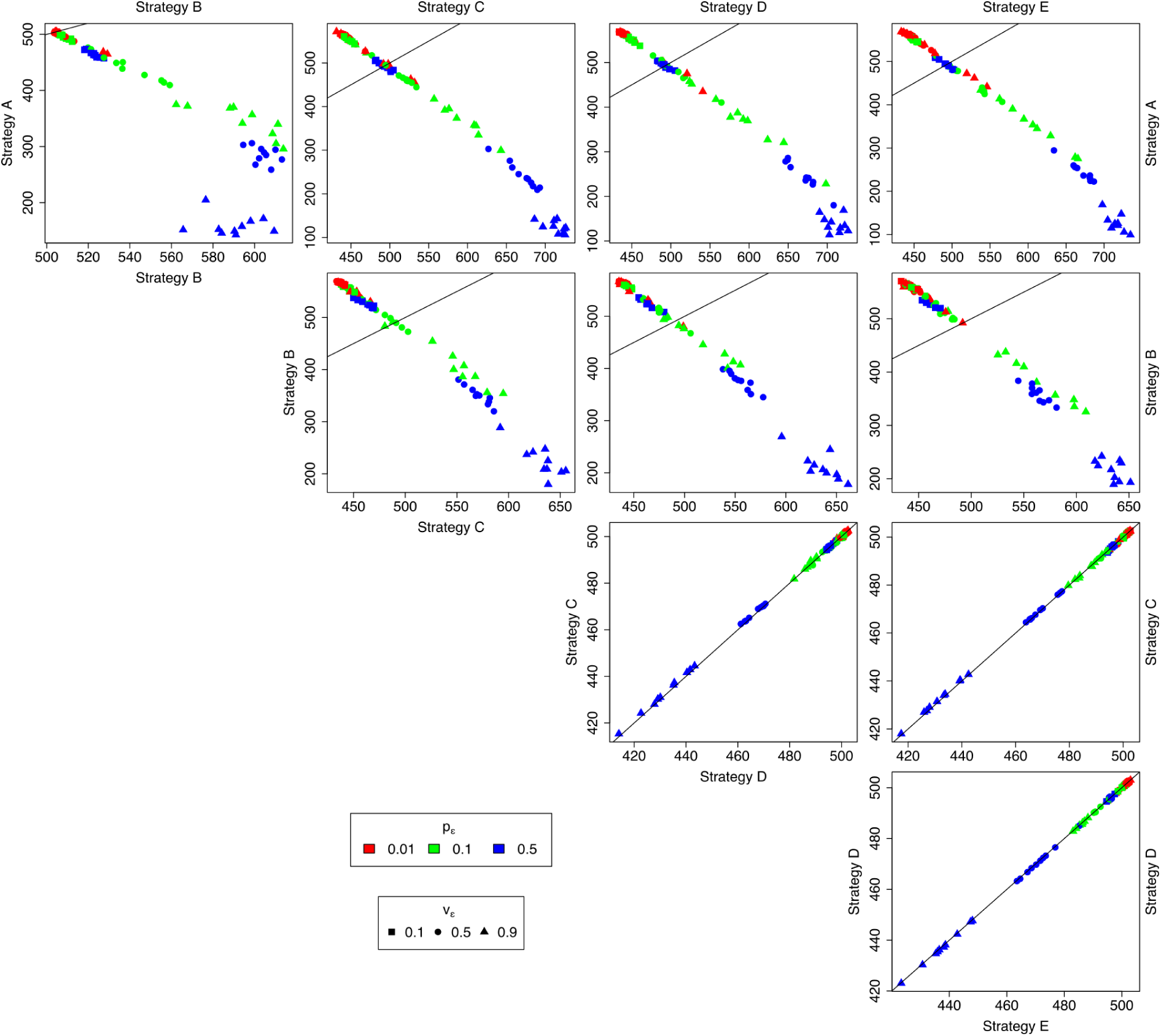
Pairwise competition between different reproductive strategies during invasion of a marginal habitat (North patch) under different probabilities of occurrence of stressful conditions 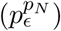 and maximum levels of stress 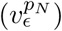, with sexual strategies exposed to a relatively weak Allee effect (*α* = 0.3 and *β* = 0.1). Axes represent the short-term temporal mean population sizes of competing strategies. Diagonal lines indicate identical temporal mean population sizes of competing reproductive strategies and therefore equivalent ecological successes. Sensitivity values: *ϕ*^*A*^ = 1.0, *ϕ*^*B*^ = 0.75, *ϕ*^*C*^ = 0.5.

**Figure S11:**
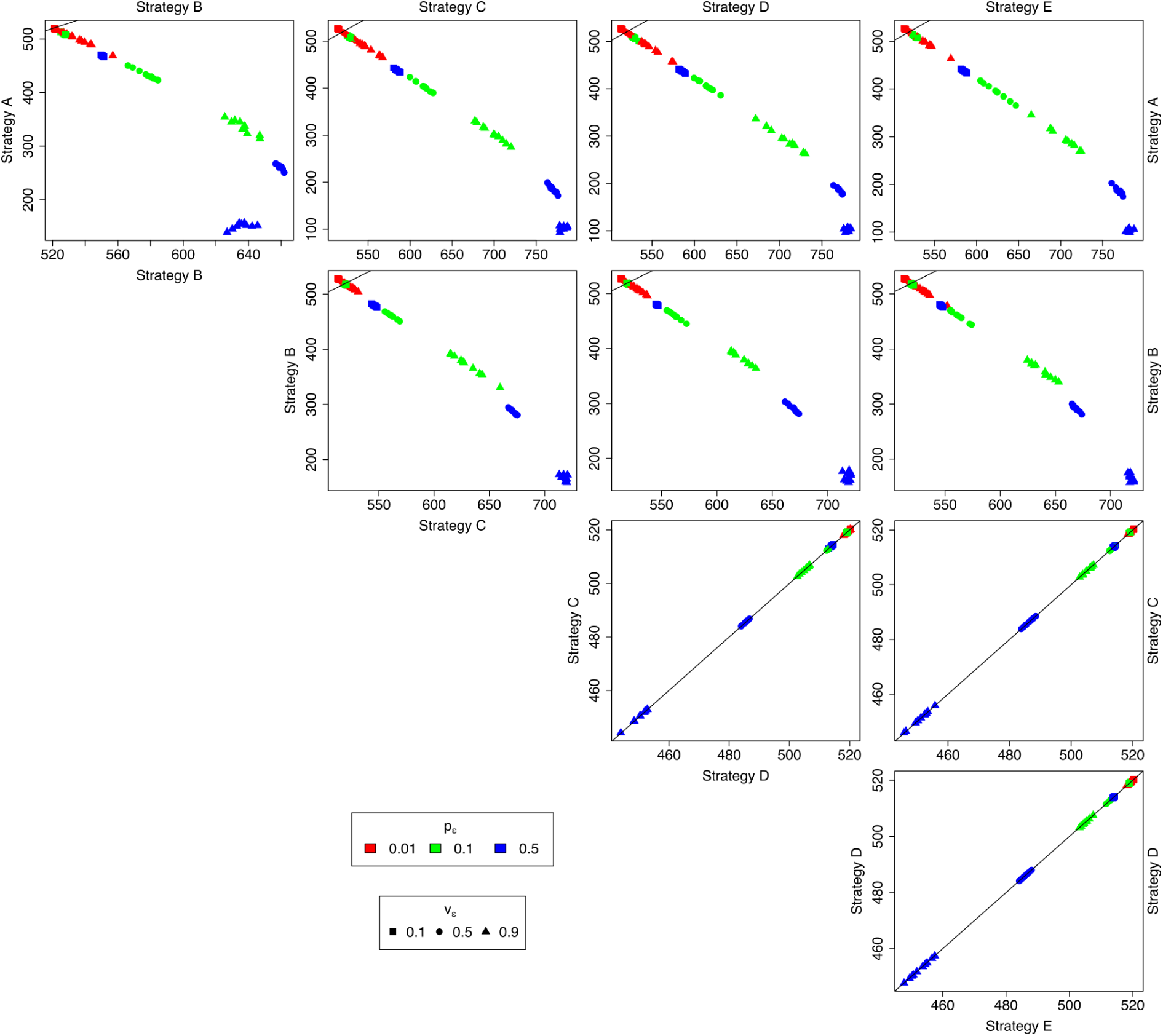
Pairwise competition between different reproductive strategies during invasion of a marginal habitat (North patch) under different probabilities of occurrence of stressful conditions 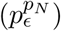 and maximum levels of stress 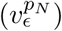, with sexual strategies exposed to a relatively weak Allee effect (*α* = 0.3 and *β* = 0.1). Axes represent the long-term temporal mean population sizes of competing strategies. Diagonal lines indicate identical temporal mean population sizes of competing reproductive strategies and therefore equivalent ecological successes. Sensitivity values: *ϕ*^*A*^ = 1.0, *ϕ*^*B*^ = 0.75, *ϕ*^*C*^ = 0.5.

**Figure S12:**
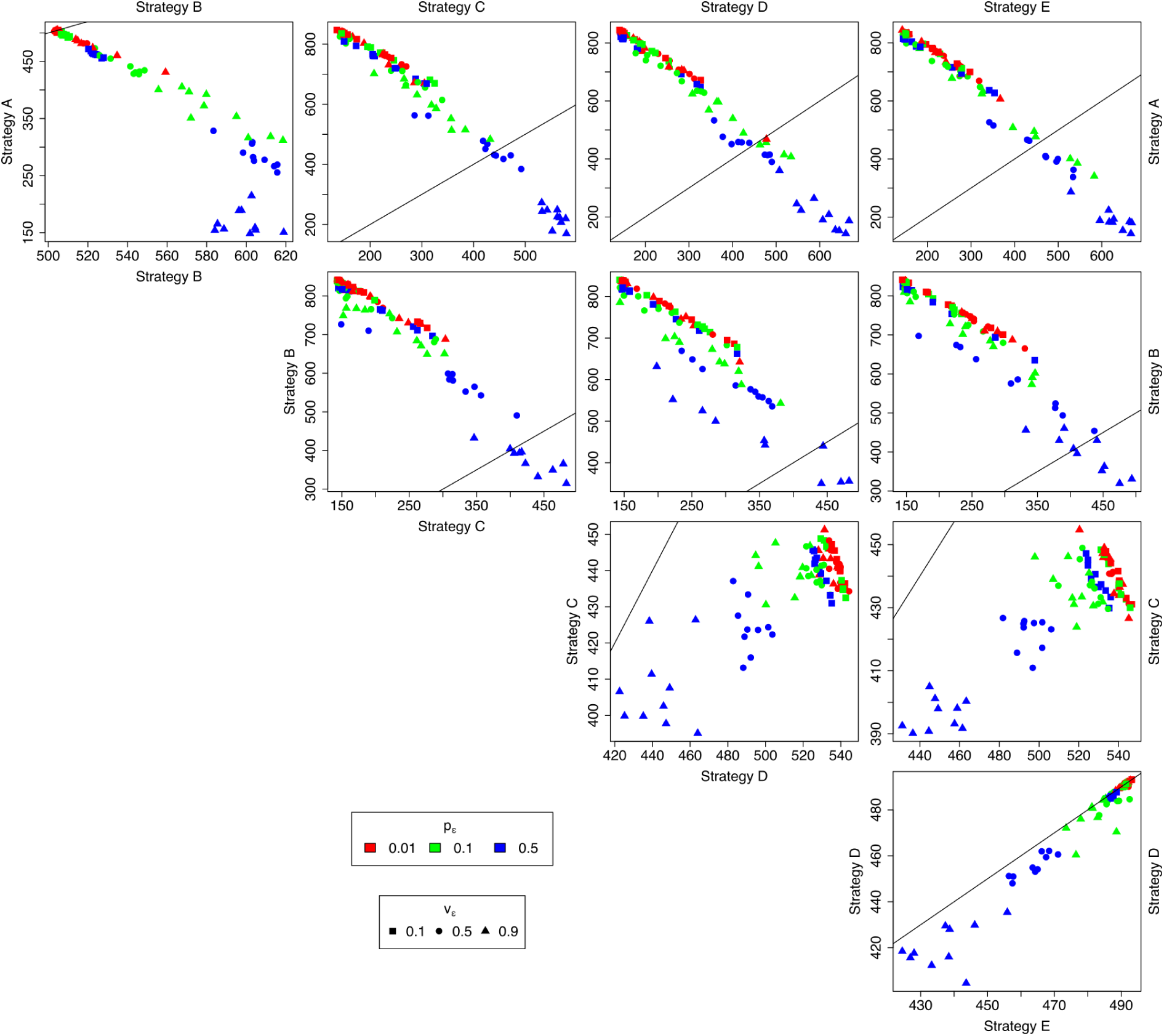
Pairwise competition between different reproductive strategies during invasion of a marginal habitat (North patch) under different probabilities of occurrence of stressful conditions 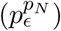 and maximum levels of stress 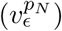, with sexual strategies exposed to a relatively strong Allee effect (*α* = 0.1 and *β* = 0.3). Axes represent the short-term temporal mean population sizes of competing strategies. Diagonal lines indicate identical temporal mean population sizes of competing reproductive strategies and therefore equivalent ecological successes. Sensitivity values: *ϕ*^*A*^ = 1.0, *ϕ*^*B*^ = 0.75, *ϕ*^*C*^ = 0.5.

**Figure S13:**
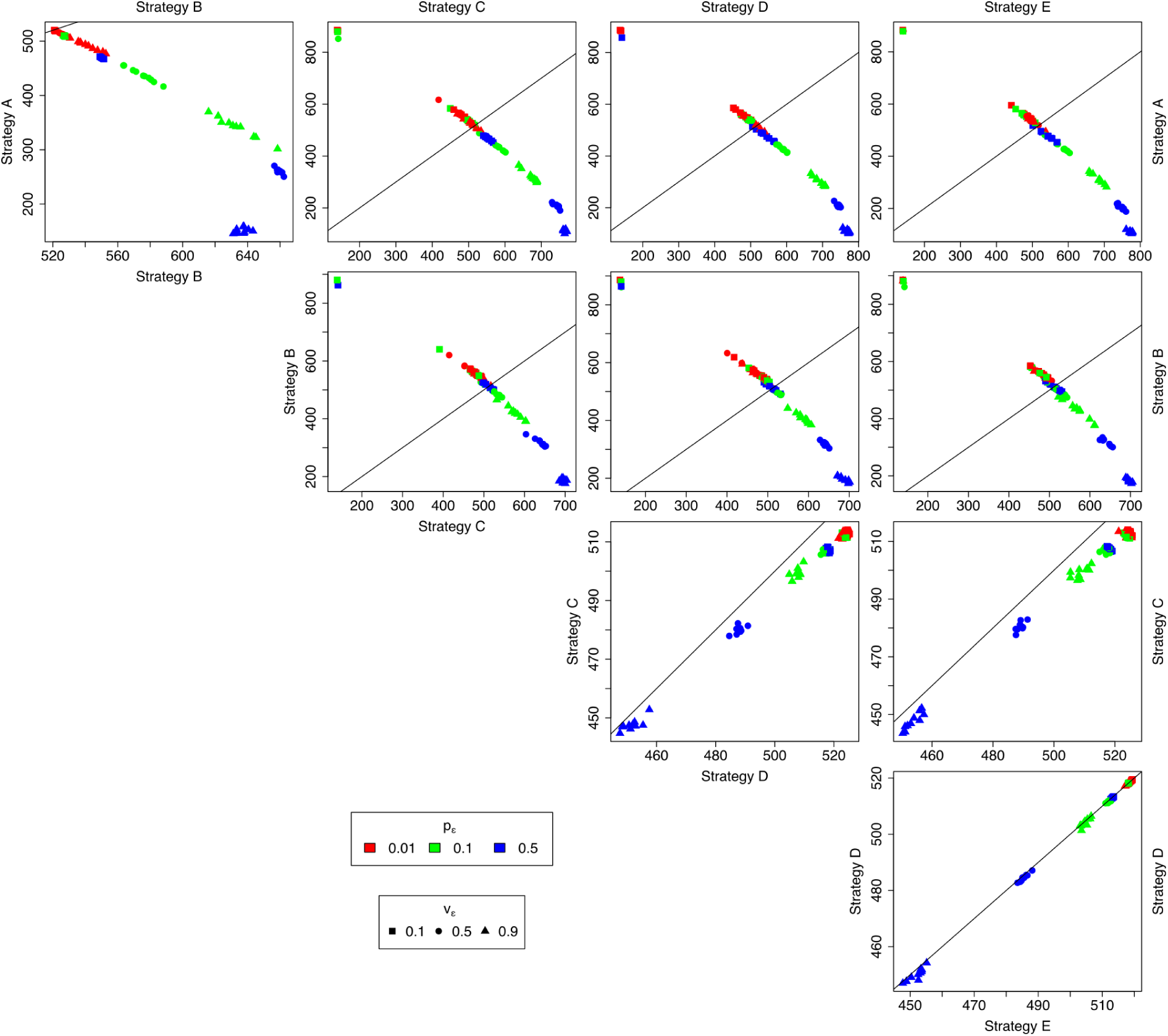
Pairwise competition between different reproductive strategies during invasion of a marginal habitat (North patch) under different probabilities of occurrence of stressful conditions 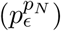 and maximum levels of stress 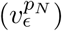, with sexual strategies exposed to a relatively strong Allee effect (*α* = 0.1 and *β* = 0.3). Axes represent the long-term temporal mean population sizes of competing strategies. Diagonal lines indicate identical temporal mean population sizes of competing reproductive strategies and therefore equivalent ecological successes. Sensitivity values: *ϕ*^*A*^ = 1.0, *ϕ*^*B*^ = 0.75, *ϕ*^*C*^ = 0.5.

**Figure S14:**
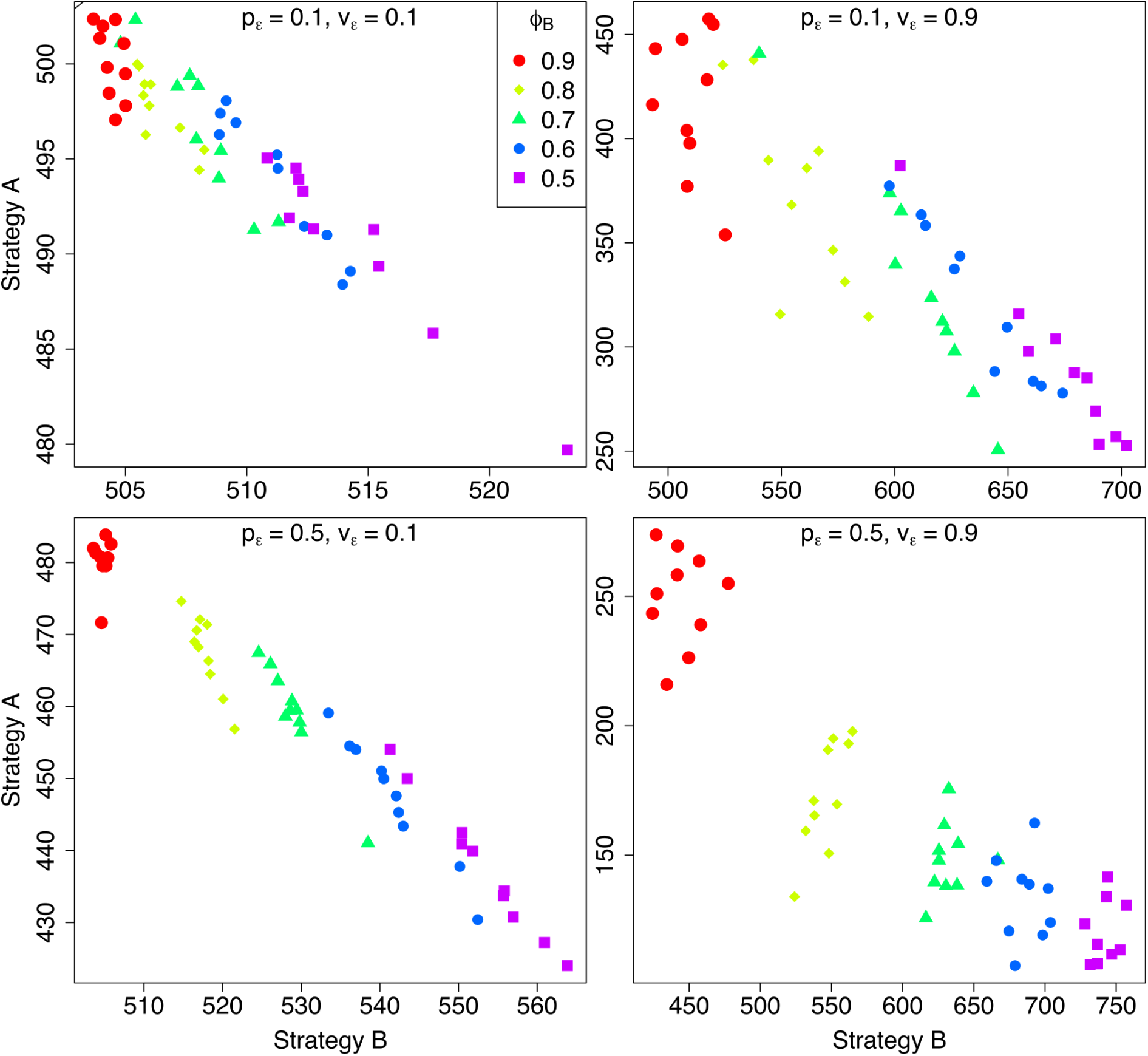
Obligate apomictic (strategy A) vs. obligate automictic parthenogenesis (strategy B). Axes represent the short-term temporal mean population sizes of competing strategies in the marginal habitat (North path). Note that *ϕ* represents the relative difference in sensitivity to environmental stress between apomictic (strategy A; *ϕ*^*A*^ = 1.0) and automictic (strategy B; *ϕ*^*B*^) parthenogenesis, or between parthenogenesis (strategy A; *ϕ*^*A*^ = 1.0) and sexual reproduction (strategy C; *ϕ*^*C*^).

**Figure S15:**
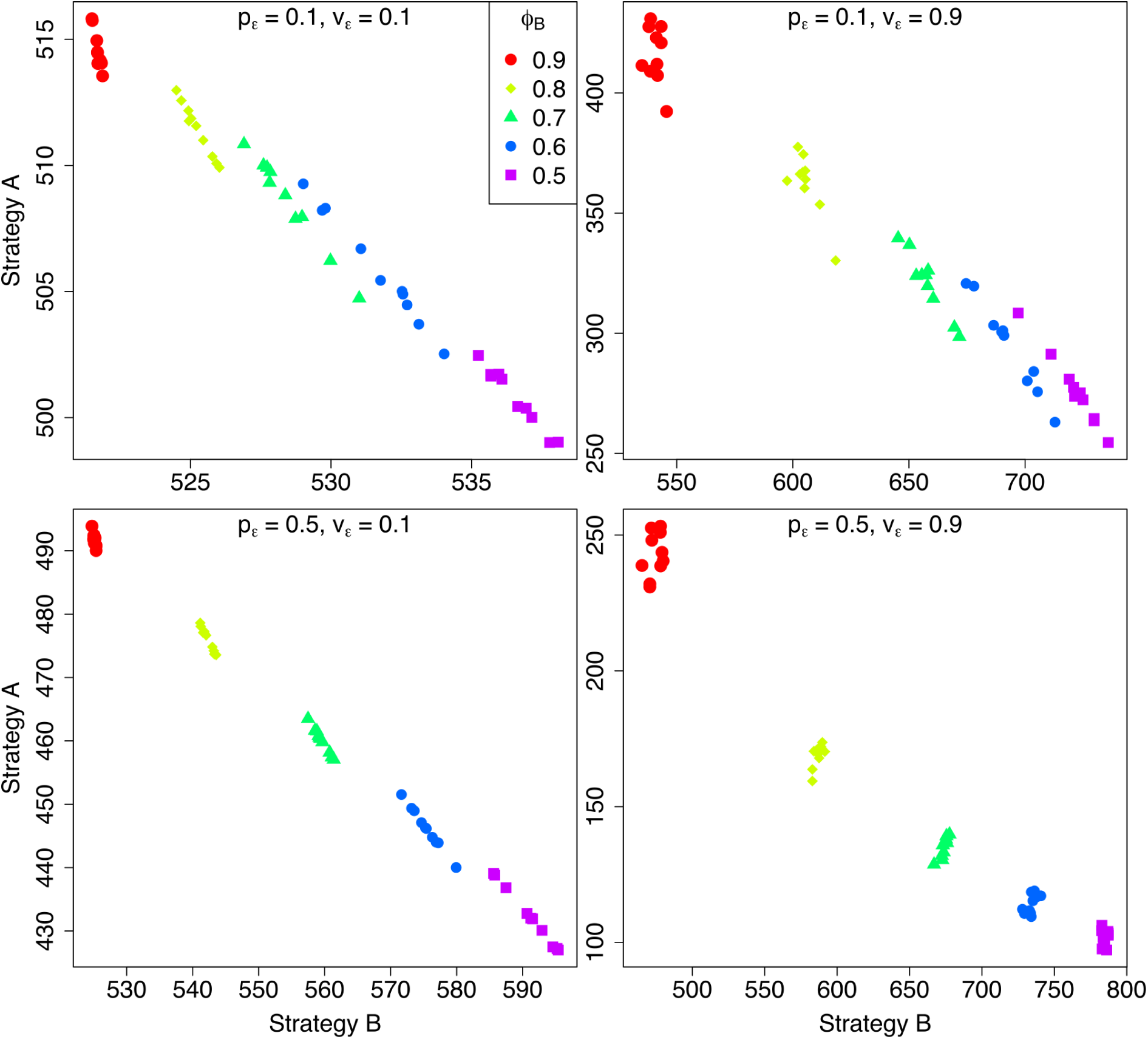
Obligate apomictic (strategy A) vs. obligate automictic parthenogenesis (strategy B). Axes represent the long-term temporal mean population sizes of competing strategies in the marginal habitat (North path). Note that *ϕ* represents the relative difference in sensitivity to environmental stress between apomictic (strategy A; *ϕ*^*A*^ = 1.0) and automictic (strategy B; *ϕ*^*B*^) parthenogenesis, or between parthenogenesis (strategy A; *ϕ*^*A*^ = 1.0) and sexual reproduction (strategy C; *ϕ*^*C*^).

**Figure S16:**
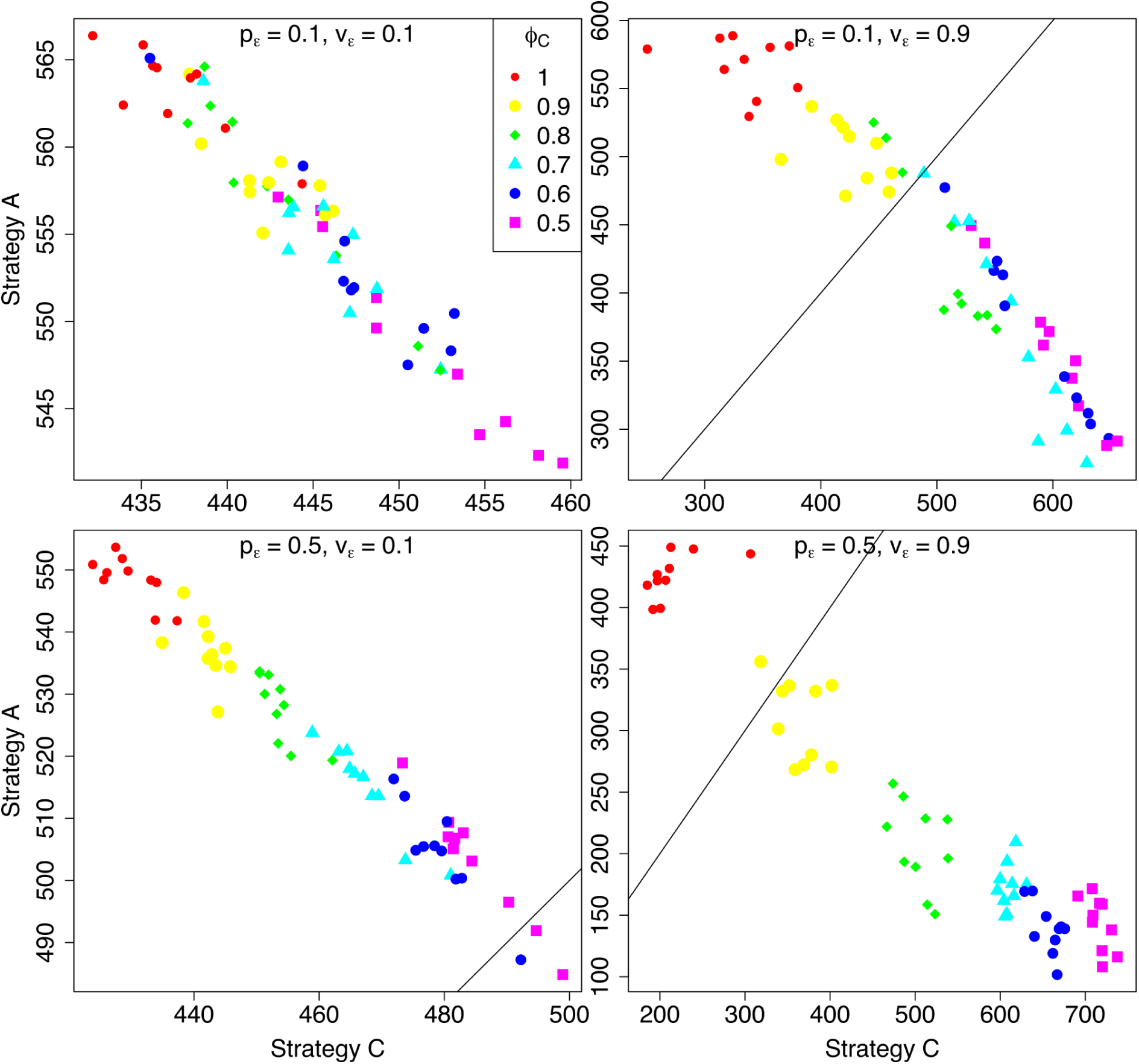
Obligate parthenogenesis (strategy A/B) vs. obligate sexual reproduction (strategy C). Axes represent the short-term temporal mean population sizes of competing strategies in the marginal habitat (North path) under a weak Allee effect (*α* = 0.3 and *β* = 0.1). Note that *ϕ* represents the relative difference in sensitivity to environmental stress between apomictic (strategy A; *ϕ*^*A*^ = 1.0) and automictic (strategy B; *ϕ*^*B*^) parthenogenesis, or between parthenogenesis (strategy A; *ϕ*^*A*^ = 1.0) and sexual reproduction (strategy C; *ϕ*^*C*^).

**Figure S17:**
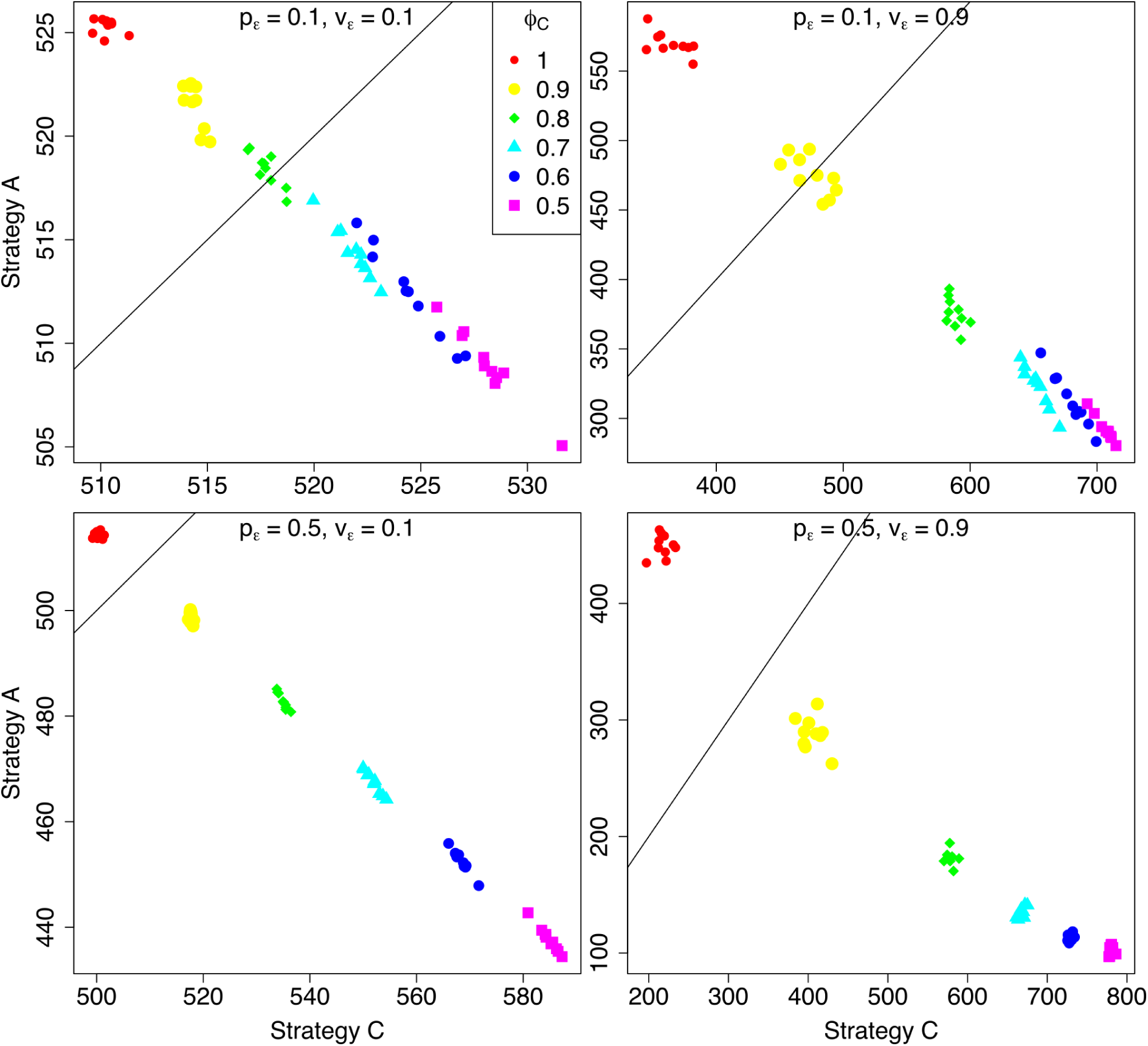
Obligate parthenogenesis (strategy A/B) vs. obligate sexual reproduction (strategy C). Axes represent the long-term temporal mean population sizes of competing strategies in the marginal habitat (North path) under a weak Allee effect (*α* = 0.3 and *β* = 0.1). Note that *ϕ* represents the relative difference in sensitivity to environmental stress between apomictic (strategy A; *ϕ*^*A*^ = 1.0) and automictic (strategy B; *ϕ*^*B*^) parthenogenesis, or between parthenogenesis (strategy A; *ϕ*^*A*^ = 1.0) and sexual reproduction (strategy C; *ϕ*^*C*^).

**Figure S18:**
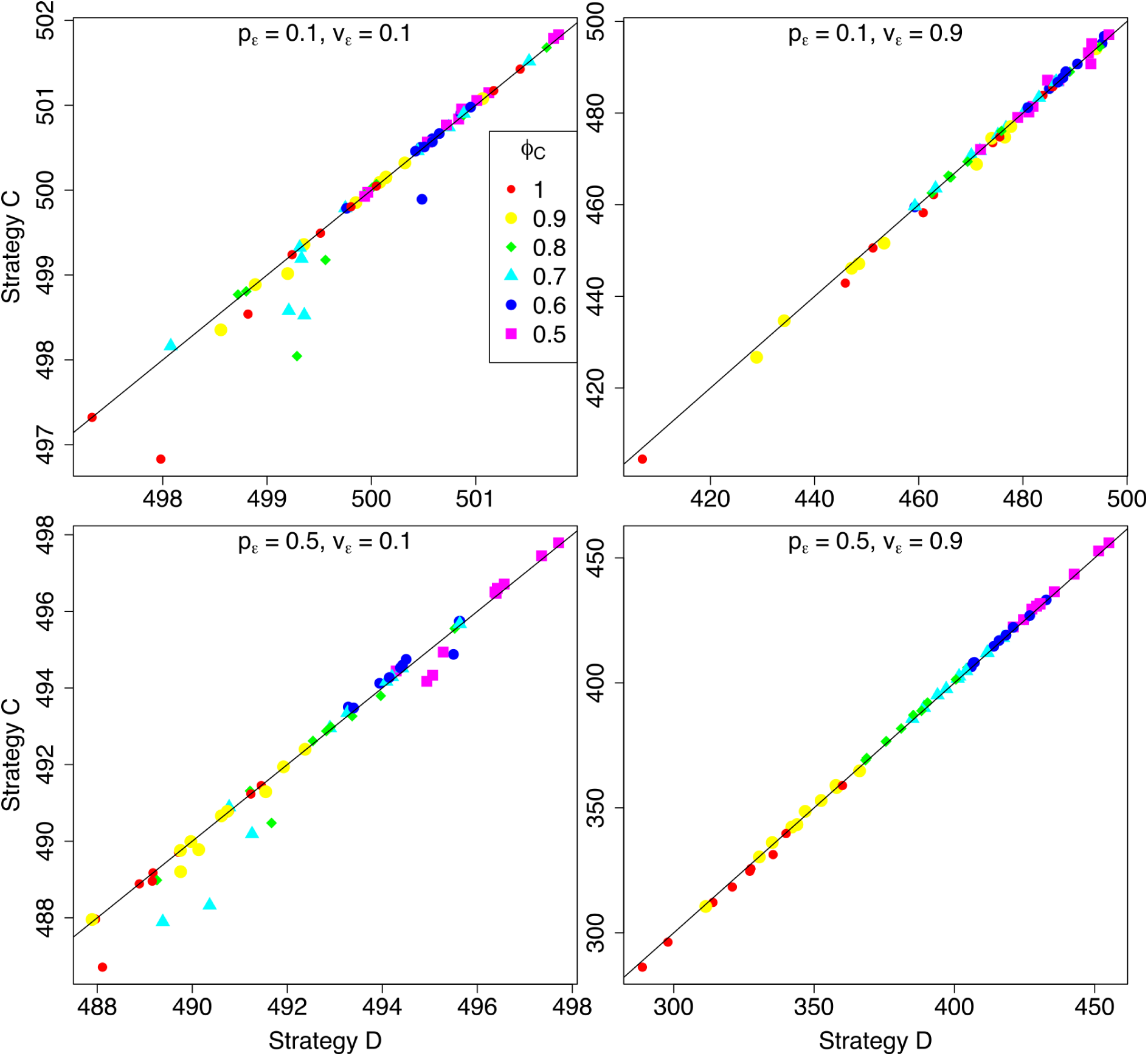
Obligate sexual reproduction (strategy C) vs. facultative parthenogenesis (strategy D/E). Axes represent the short-term temporal mean population sizes of competing strategies in the marginal habitat (North path) under a weak Allee effect (*α* = 0.3 and *β* = 0.1). Note that *ϕ* represents the relative difference in sensitivity to environmental stress between apomictic (strategy A; *ϕ*^*A*^ = 1.0) and automictic (strategy B; *ϕ*^*B*^) parthenogenesis, or between parthenogenesis (strategy A; *ϕ*^*A*^ = 1.0) and sexual reproduction (strategy C; *ϕ*^*C*^).

**Figure S19:**
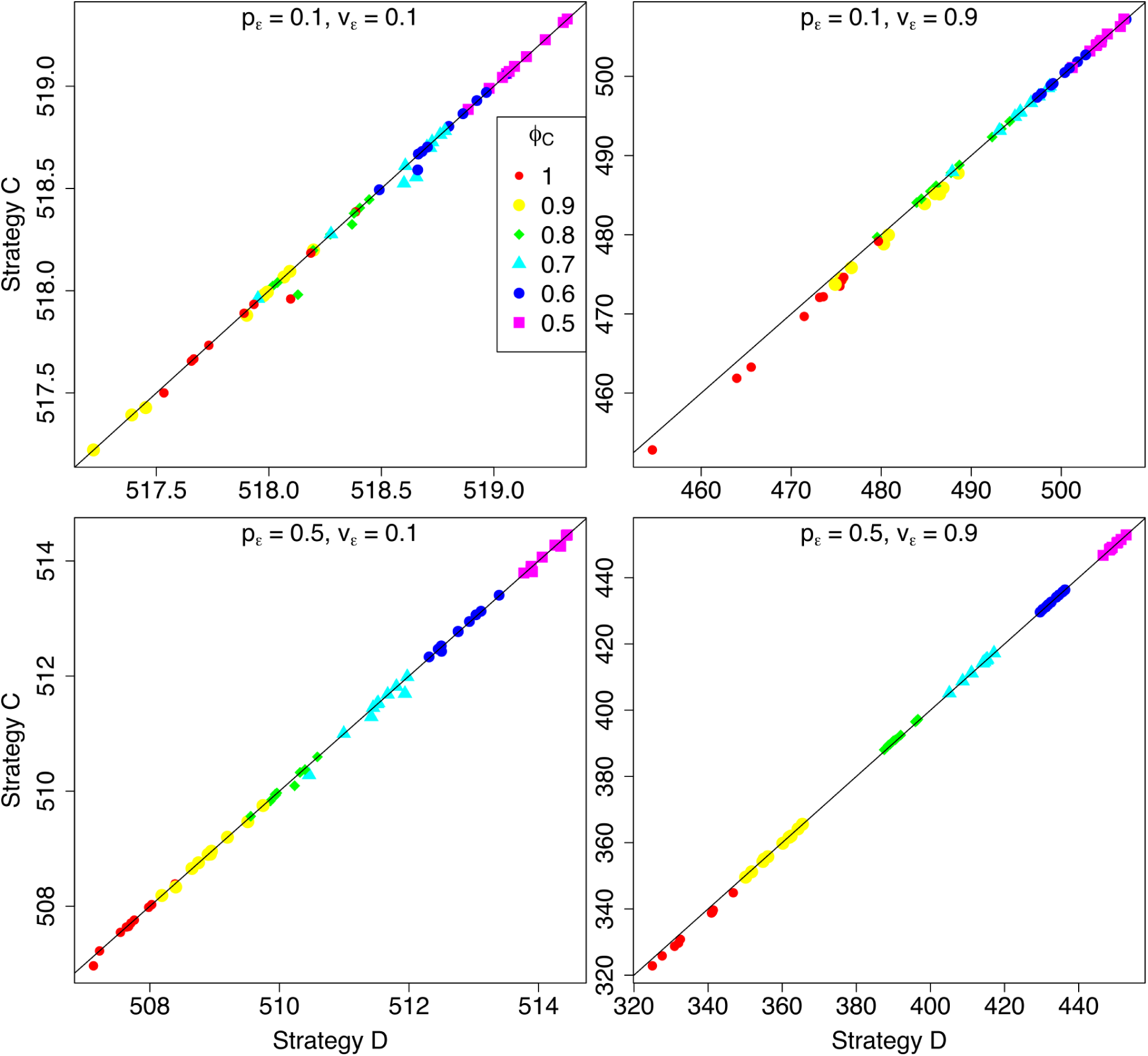
Obligate sexual reproduction (strategy C) vs. facultative parthenogenesis (strategy D/E). Axes represent the long-term temporal mean population sizes of competing strategies in the marginal habitat (North path) under a weak Allee effect (*α* = 0.3 and *β* = 0.1). Note that *ϕ* represents the relative difference in sensitivity to environmental stress between apomictic (strategy A; *ϕ*^*A*^ = 1.0) and automictic (strategy B; *ϕ*^*B*^) parthenogenesis, or between parthenogenesis (strategy A; *ϕ*^*A*^ = 1.0) and sexual reproduction (strategy C; *ϕ*^*C*^).

**Figure S20:**
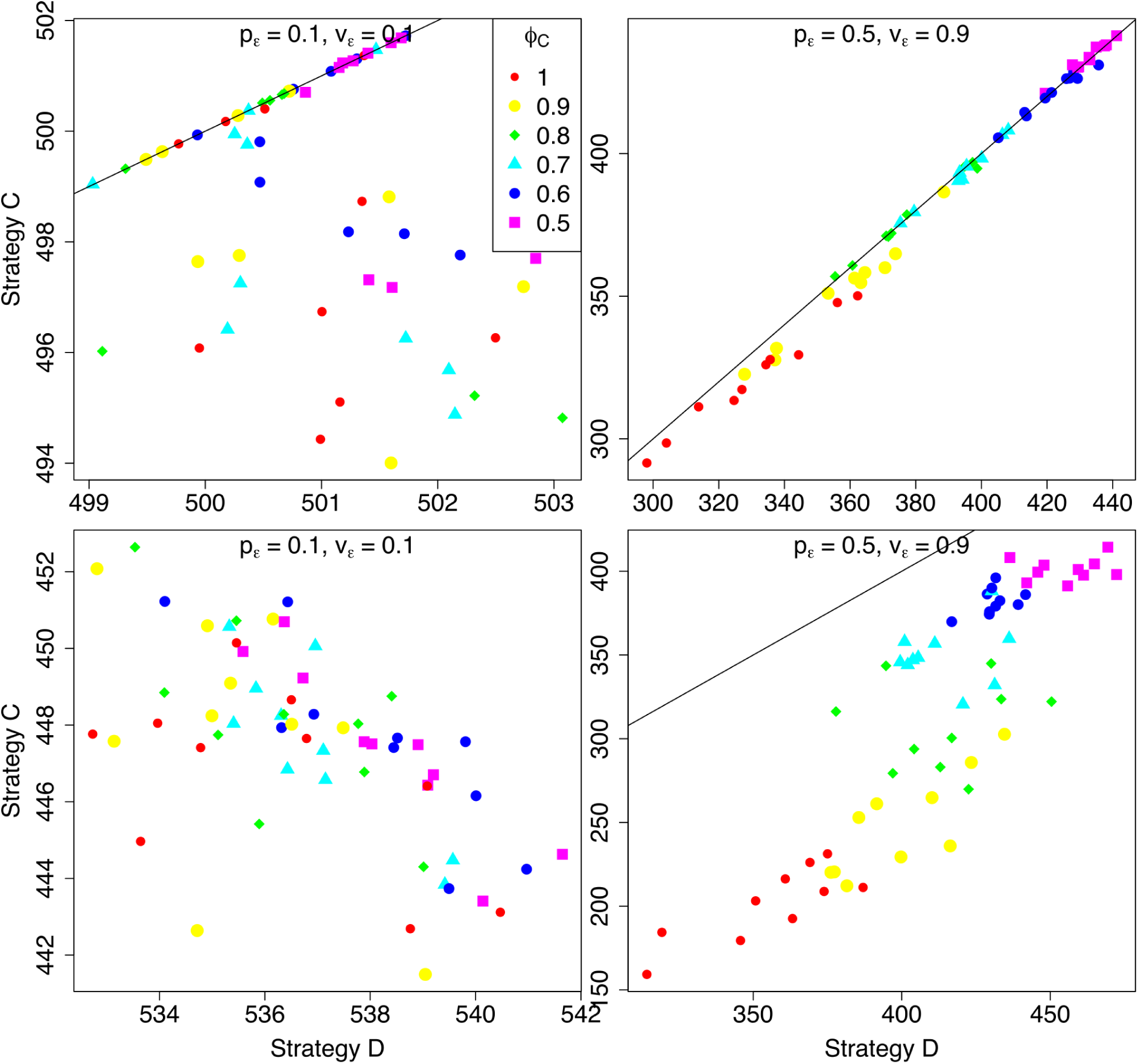
Obligate sexual reproduction (strategy C) vs. facultative parthenogenesis (strategy D/E). Axes represent the short-term temporal mean population sizes of competing strategies in the marginal habitat (North path) under a weak Allee effect (top row; *α* = 0.3 and *β* = 0.1) and a strong Allee effect (bottom row; *α* = 0.1 and *β* = 0.3), with a high transition rate (*τ* = 0.8). Note that *ϕ* represents the relative difference in sensitivity to environmental stress between apomictic (strategy A; *ϕ*^*A*^ = 1.0) and automictic (strategy B; *ϕ*^*B*^) parthenogenesis, or between parthenogenesis (strategy A; *ϕ*^*A*^ = 1.0) and sexual reproduction (strategy C; *ϕ*^*C*^). high transition rate (*τ* = 0.8).

**Figure S21:**
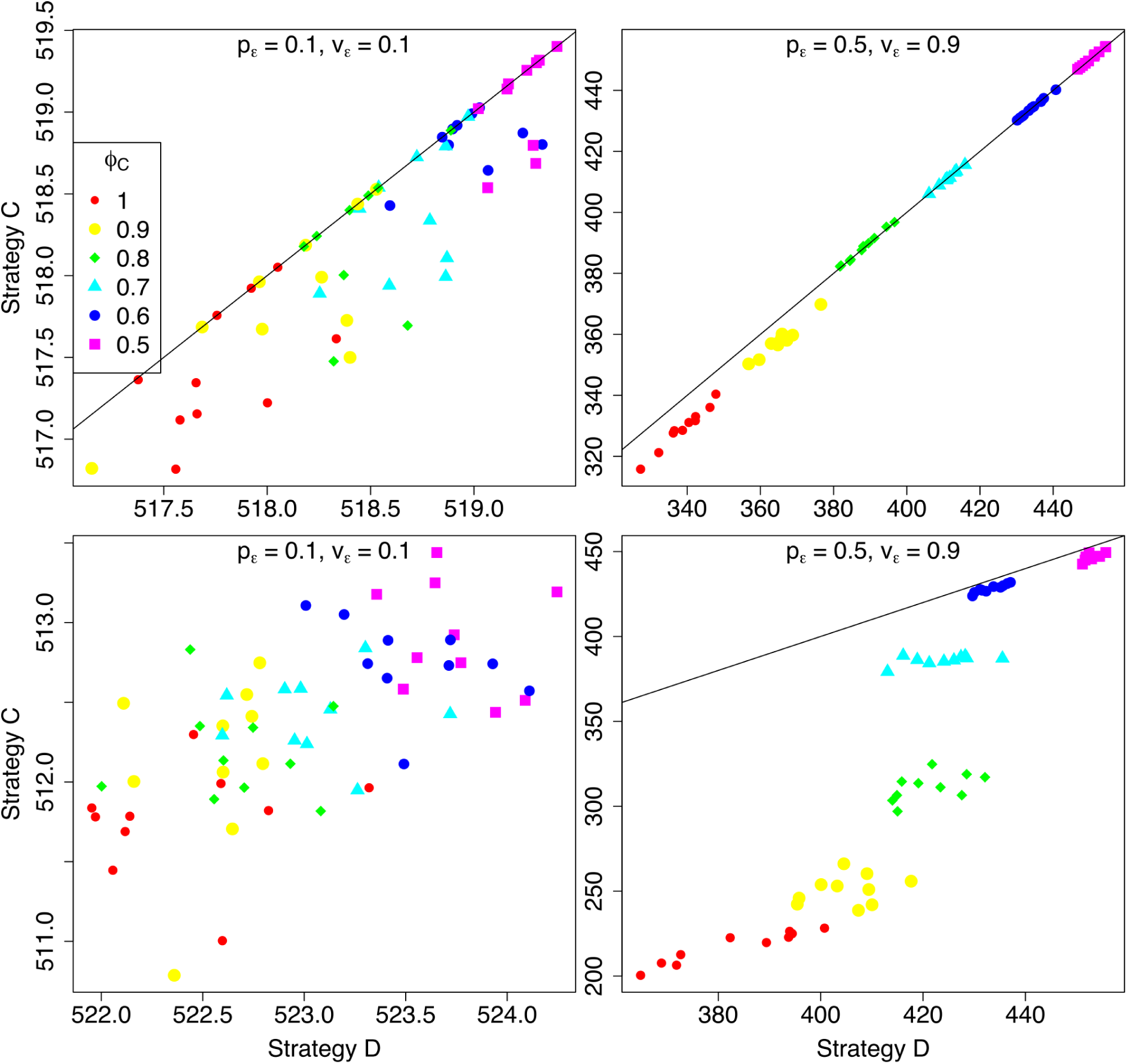
Obligate sexual reproduction (strategy C) vs. facultative parthenogenesis (strategy D/E). Axes represent the long-term temporal mean population sizes of competing strategies in the marginal habitat (North path) under a weak Allee effect (top row; *α* = 0.3 and *β* = 0.1) and a strong Allee effect (bottom row; *α* = 0.1 and *β* = 0.3), with a high transition rate (*τ* = 0.8). Note that *ϕ* represents the relative difference in sensitivity to environmental stress between apomictic (strategy A; *ϕ*^*A*^ = 1.0) and automictic (strategy B; *ϕ*^*B*^) parthenogenesis, or between parthenogenesis (strategy A; *ϕ*^*A*^ = 1.0) and sexual reproduction (strategy C; *ϕ*^*C*^). high transition rate (*τ* = 0.8)..

**Figure S22:**
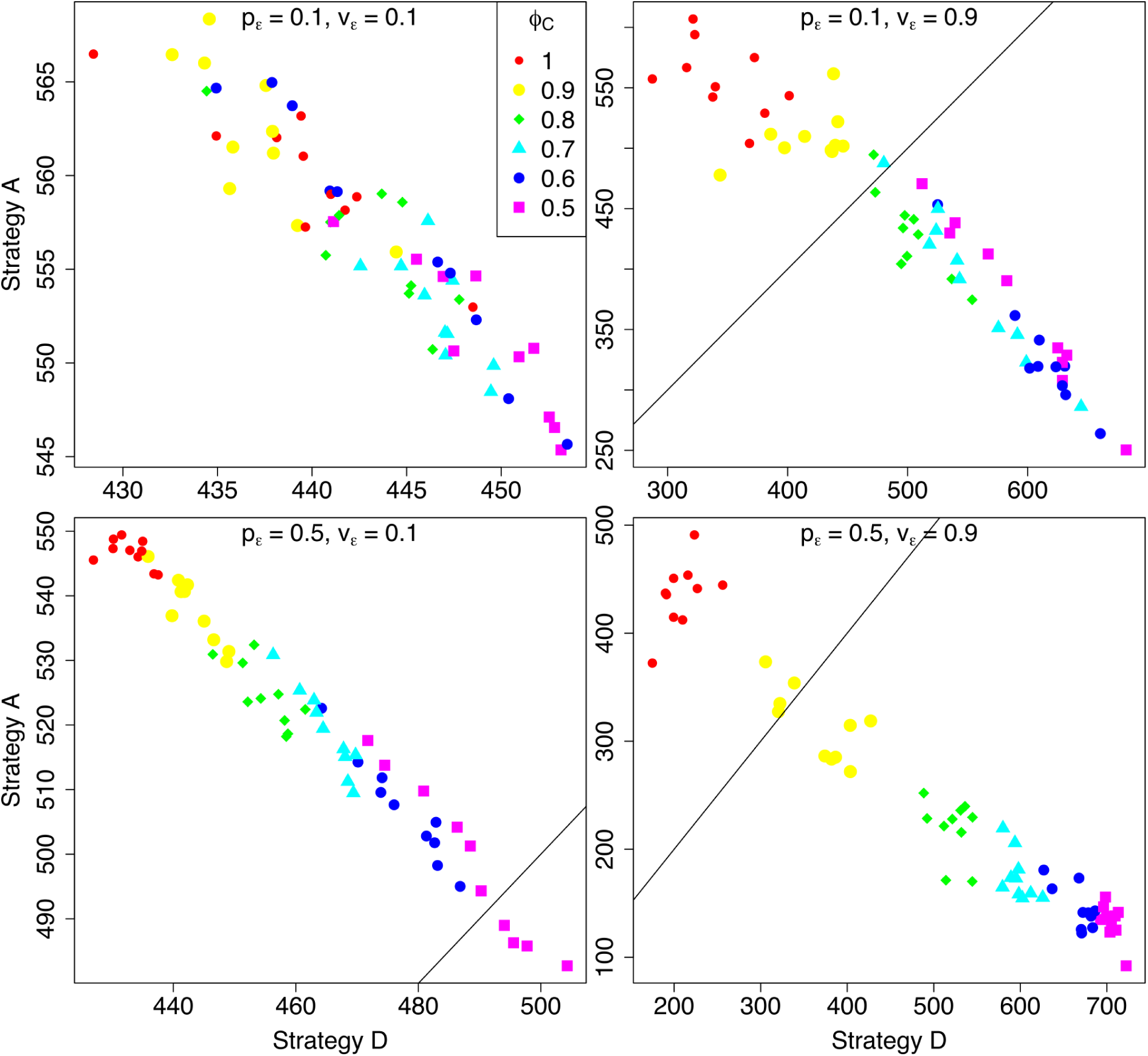
Obligate parthenogenesis (strategy A/B) vs. facultative parthenogenesis (strategy D/E). Axes represent the short-term temporal mean population sizes of competing strategies in the marginal habitat (North path) under a weak Allee effect (*α* = 0.3 and *β* = 0.1). Note that *ϕ* represents the relative difference in sensitivity to environmental stress between apomictic (strategy A; *ϕ*^*A*^ = 1.0) and automictic (strategy B; *ϕ*^*B*^) parthenogenesis, or between parthenogenesis (strategy A; *ϕ*^*A*^ = 1.0) and sexual reproduction (strategy C; *ϕ*^*C*^).

**Figure S23:**
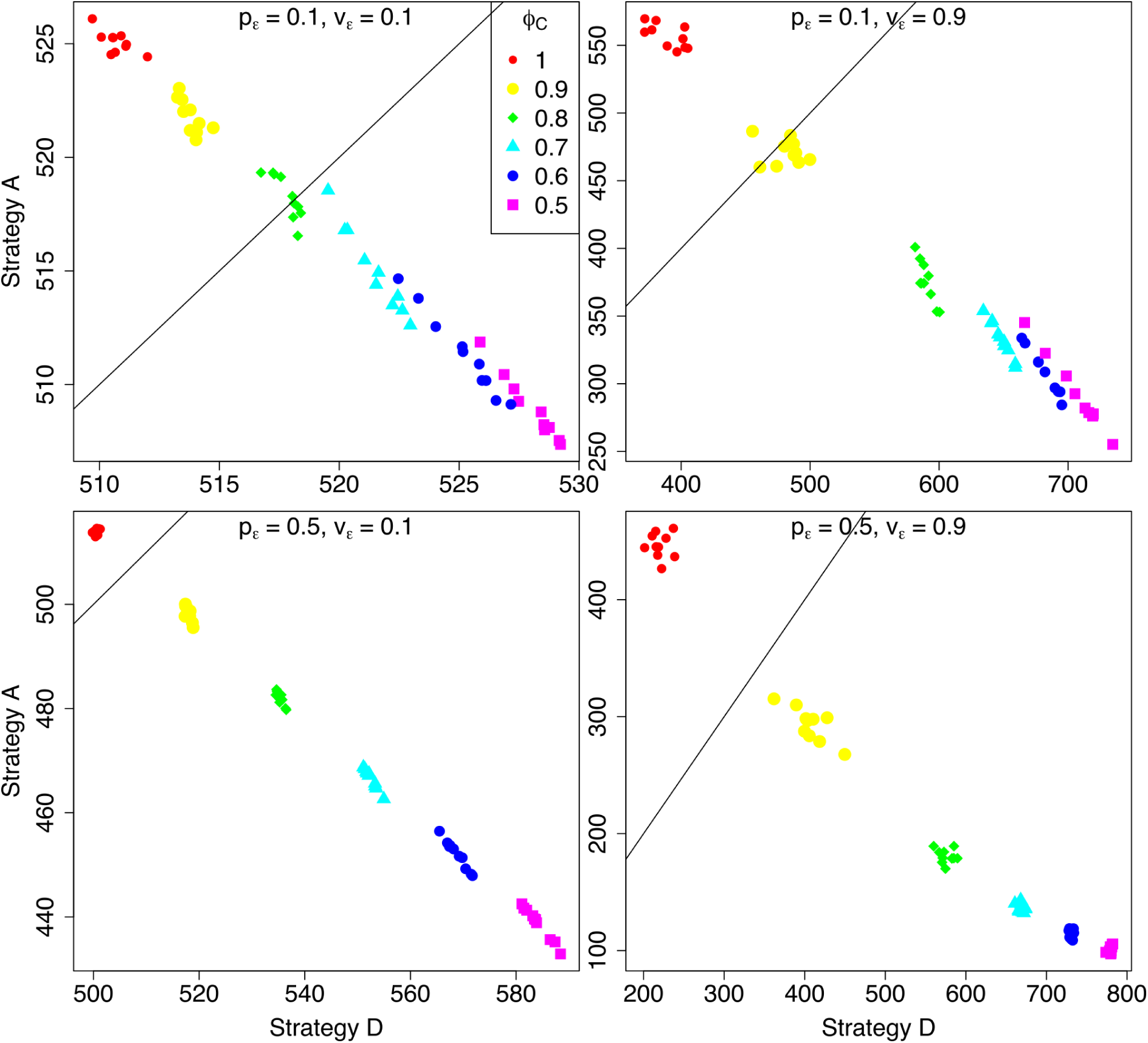
Obligate parthenogenesis (strategy A/B) vs. facultative parthenogenesis (strategy D/E). Axes represent the long-term temporal mean population sizes of competing strategies in the marginal habitat (North path) under a weak Allee effect (*α* = 0.3 and *β* = 0.1). Note that *ϕ* represents the relative difference in sensitivity to environmental stress between apomictic (strategy A; *ϕ*^*A*^ = 1.0) and automictic (strategy B; *ϕ*^*B*^) parthenogenesis, or between parthenogenesis (strategy A; *ϕ*^*A*^ = 1.0) and sexual reproduction (strategy C; *ϕ*^*C*^).

**Figure S24:**
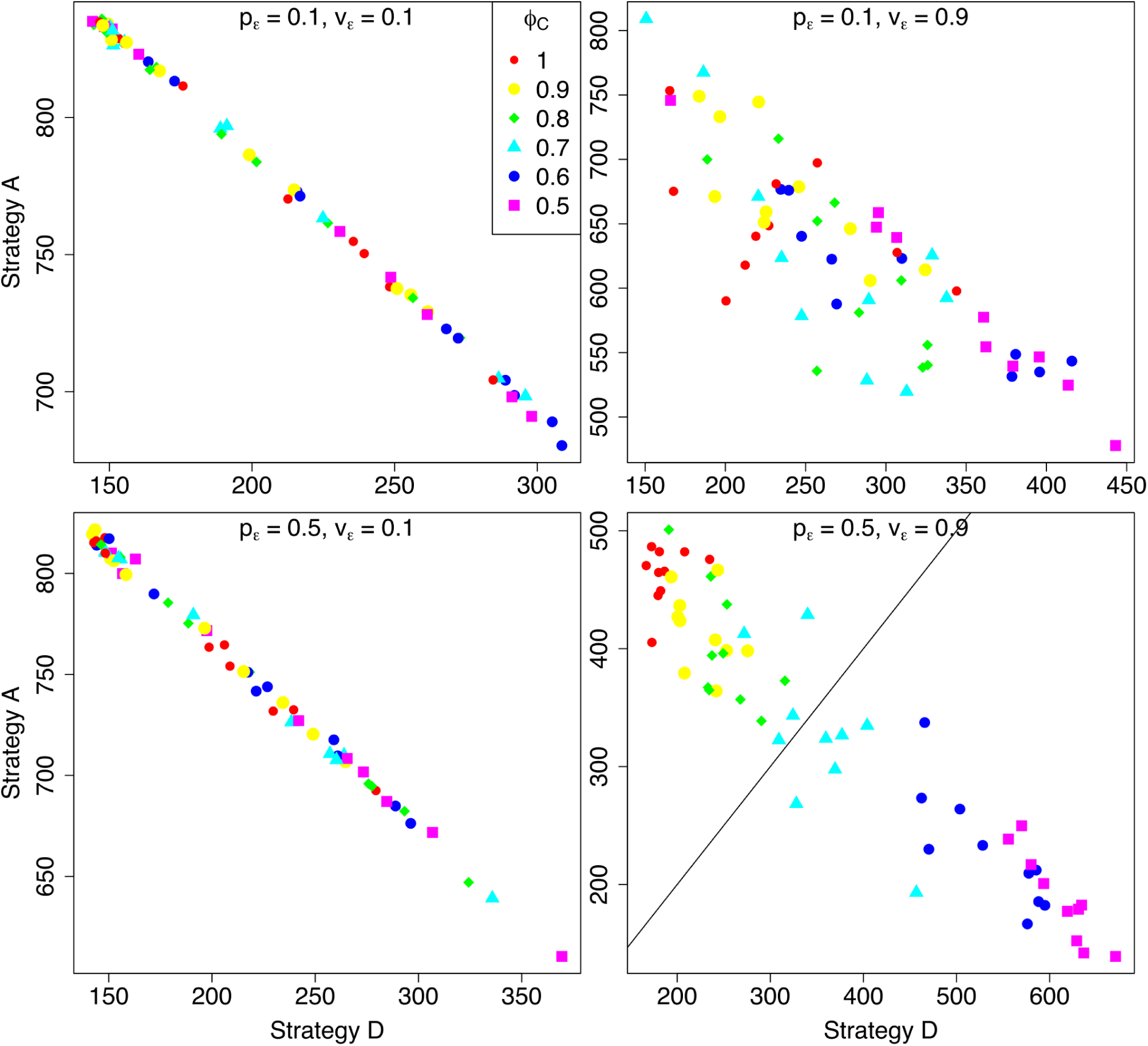
Obligate parthenogenesis (strategy A/B) vs. facultative parthenogenesis (strategy D/E). Axes represent the short-term temporal mean population sizes of competing strategies in the marginal habitat (North path) under a strong Allee effect (*α* = 0.1 and *β* = 0.3). Note that *ϕ* represents the relative difference in sensitivity to environmental stress between apomictic (strategy A; *ϕ*^*A*^ = 1.0) and automictic (strategy B; *ϕ*^*B*^) parthenogenesis, or between parthenogenesis (strategy A; *ϕ*^*A*^ = 1.0) and sexual reproduction (strategy C; *ϕ*^*C*^).

**Figure S25:**
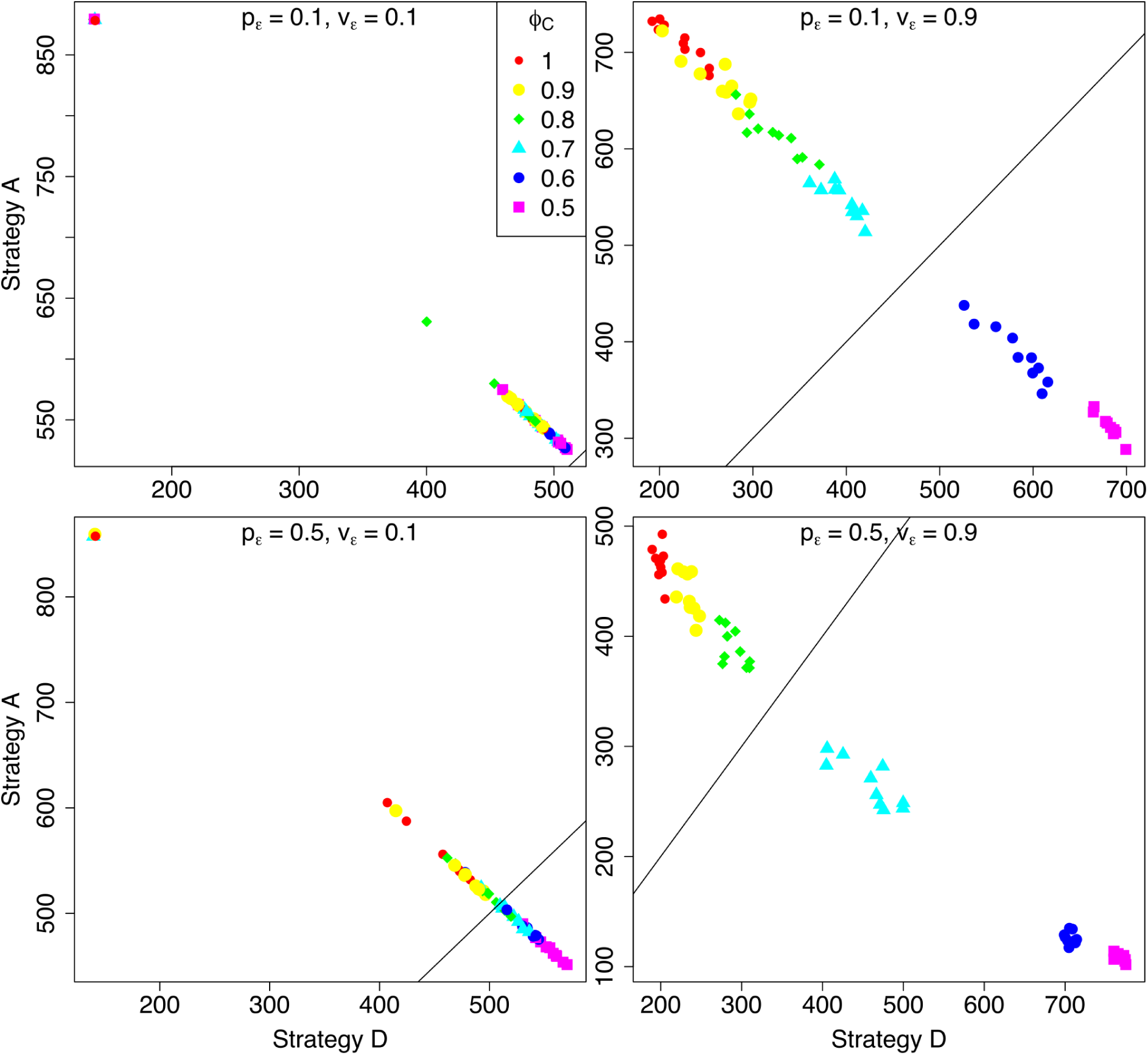
Obligate parthenogenesis (strategy A/B) vs. facultative parthenogenesis (strategy D/E). Axes represent the long-term temporal mean population sizes of competing strategies in the marginal habitat (North path) under a strong Allee effect (*α* = 0.1 and *β* = 0.3). Note that *ϕ* represents the relative difference in sensitivity to environmental stress between apomictic (strategy A; *ϕ*^*A*^ = 1.0) and automictic (strategy B; *ϕ*^*B*^) parthenogenesis, or between parthenogenesis (strategy A; *ϕ*^*A*^ = 1.0) and sexual reproduction (strategy C; *ϕ*^*C*^).

## Literature Cited

[1] Tilquin A, Kokko H. What does the geography of parthenogenesis teach us about sex? Philosophical Transactions of the Royal Society B: Biological Sciences. 2016;371(1706):20150538. doi: 10.1098/rstb.2015.0538.

[2] Dawson KJ. The advantage of asexual reproduction: when is it two-fold? Journal of Theoretical Biology. 1995;176(3):341–347. doi: 10.1006/jtbi.1995.0203.

[3] de Meeûs T, Prugnolle F, Agnew P. Asexual reproduction: genetics and evolutionary aspects. Cellular and Molecular Life Sciences. 2007;64(11):1355–1372. doi: 10.1007/s00018-007-6515-2.

[4] Williams GC, Mitton JB. Why reproduce sexually? Journal of Theoretical Biology. 1973;39(3):545–554. doi: 10.1016/0022-5193(73)90067-2.

[5] Crow JF. Advantages of sexual reproduction. Developmental Genetics. 1994;15(3):205–213. doi: 10.1002/dvg.1020150303.

[6] Crow JF. An advantage of sexual reproduction in a rapidly changing environment. Journal of Heredity. 1992;83(3):169–173. doi: 10.1093/oxfordjournals.jhered.a111187.

[7] Bengtsson BO. Genetic variation in organisms with sexual and asexual reproduction. Journal of Evolutionary Biology. 2003;16(2):189–199. doi: 10.1046/j.1420-9101.2003.00523.x.

[8] Kellner K, Heinze J. Mechanism of facultative parthenogenesis in the ant *Platythyrea punctata*. Evolutionary Ecology. 2011;25(1):77–89. doi: 10.1007/s10682-010-9382-5.

[9] Chang CC, Ting CT, Chang CH, Fang S, Chang HY. The persistence of facultative parthenogenesis in *Drosophila albomicans*. PLoS ONE. 2014;9(11):e113275. doi: 10.1371/journal.pone.0113275.

[10] Fournier D, Hellemans S, Hanus R, Roisin Y. Facultative asexual reproduction and genetic diversity of populations in the humivorous termite *Cavitermes tuberosus*. Proceedings of the Royal Society B: Biological Sciences. 2016;283(1832):20160196. doi: 10.1098/rspb.2016.0196.

[11] Dudgeon CL, Coulton L, Bone R, Ovenden JR, Thomas S. Switch from sexual to parthenogenetic reproduction in a zebra shark. Scientific Reports. 2017;7:40537. doi: 10.1038/srep40537.

[12] Vandel A. La parthénogenèse géographique. Contribution à l’étude biologique et cytologique de la parthénogenèse naturelle. Bulletin biologique de la France et de la Belgique. 1928;62:164–281.

[13] Stieha C, García-Ramos G, Nicholas McLetchie D, Crowley P. Maintenance of the sexes and persistence of a clonal organism in spatially complex metapopulations. Evolutionary Ecology. 2017;31(3):363–386. doi: 10.1007/s10682-016-9841-8.

[14] Glesener RR, Tilman D. Sexuality and the components of environmental uncertainty: clues from geographic parthenogenesis in terrestrial animals. The American Naturalist. 1978;112(986):659–673. doi: 10.1086/283308.

[15] Bierzychudek P. Patterns in plant parthenogenesis. Experientia. 1985;41(10):1255–1264. doi: 10.1007/BF01952068.

[16] Haag CR, Ebert D. A new hypothesis to explain geographic parthenogenesis. Annales Zoologici Fennici. 2004;41(4):539–544.

[17] Johnson SG. Geographic ranges, population structure, and ages of sexual and parthenogenetic snail lineages. Evolution. 2006;60(7):1417–1426. doi: 10.1111/j.0014-3820.2006.tb01220.x.

[18] Hörandl E. The complex causality of geographical parthenogenesis. New Phytologist. 2006;171(3):525–538. doi: 10.1111/j.1469-8137.2006.01769.x.

[19] Kearney M. Hybridization, glaciation and geographical parthenogenesis. Trends in Ecology & Evolution. 2005;20(9):495–502. doi: 10.1016/j.tree.2005.06.005.

[20] Peck JR, Yearsley JM, Waxman D. Explaining the geographic distributions of sexual and asexual populations. Nature. 1998;391(6670):889–892. doi: 10.1038/36099.

[21] Law JH, Crespi BJ. The evolution of geographic parthenogenesis in *Timema* walking-sticks. Molecular Ecology. 2002;11(8):1471–1489. doi: 10.1046/j.1365-294x.2002.01547.x.

[22] Hiraoka M, Higa M. Novel distribution pattern between coexisting sexual and obligate asexual variants of the true estuarine macroalga *Ulva prolifera*. Ecology and Evolution. 2016;6(11):3658–3671. doi: 10.1002/ece3.2149.

[23] Gabrielsen TM, Brochmann C, Rueness J. The Baltic Sea as a model system for studying postglacial colonization and ecological differentiation, exemplified by the red alga *Ceramium tenuicorne*. Molecular Ecology. 2002;11(10):2083–2095. doi: 10.1046/j.1365-294X.2002.01601.x.

[24] Tatarenkov A, Bergström L, Jönsson RB, Serrão EA, Kautsky L, Johannesson K. Intriguing asexual life in marginal populations of the brown seaweed *Fucus vesiculosus*. Molecular Ecology. 2005;14(2):647–651. doi: 10.1111/j.1365-294X.2005.02425.x.

[25] Pereyra RT, Bergström L, Kautsky L, Johannesson K. Rapid speciation in a newly opened postglacial marine environment, the Baltic Sea. BMC Evolutionary Biology. 2009;9:70. doi: 10.1186/1471-2148-9-70.

[26] Johannesson K, Johansson D, Larsson KH, Huenchuñir CJ, Perus J, Forslund H, et al. Frequent clonality in fucoids (*Fucus radicans* and *Fucus vesiculosus*; Fucales, Phaeophyceae) in the Baltic Sea. Journal of Phycology. 2011;47(5):990–998. doi: 10.1111/j.1529-8817.2011.01032.x.

[27] Rafajlović M, Kleinhans D, Gulliksson C, Fries J, Johansson D, Ardehed A, et al. Neutral processes forming large clones during colonization of new areas. Journal of Evolutionary Biology. 2017;30(8):1544–1560. doi: 10.1111/jeb.13124.

[28] Hojsgaard D, Hörandl E. The rise of apomixis in natural plant populations. Frontiers in Plant Science. 2019;10. doi: 10.3389/fpls.2019.00358.

[29] Kazuki S, Tojo K. Automictic parthenogenesis of a geographically parthenogenetic mayfly, *Ephoron shigae* (Insecta: Ephemeroptera, Polymitarcyidae). Biological Journal of the Linnean Society. 2010;99(2):335–343. doi: 10.1111/j.1095-8312.2009.01351.x.

[30] Cosendai AC, Wagner J, Ladinig U, Rosche C, Hörandl E. Geographical parthenogenesis and population genetic structure in the alpine species *Ranunculus kuepferi* (Ranunculaceae). Heredity. 2013;110(6):560–569. doi: 10.1038/hdy.2013.1.

[31] Menken SBJ, Smit E, Nijs HJCMD. Genetical population structure in plants: gene flow between diploid sexual and triploid asexual dandelions (*Taraxacum* section Ruderalia). Evolution; International Journal of Organic Evolution. 1995;49(6):1108–1118. doi: 10.1111/j.1558-5646.1995.tb04437.x.

[32] Dijk PJv. Ecological and evolutionary opportunities of apomixis: insights from *Taraxacum* and *Chondrilla*. Philosophical Transactions of the Royal Society of London Series B: Biological Sciences. 2003;358(1434):1113–1121. doi: 10.1098/rstb.2003.1302.

[33] Tucker AE, Ackerman MS, Eads BD, Xu S, Lynch M. Population-genomic insights into the evolutionary origin and fate of obligately asexual *Daphnia pulex*. Proceedings of the National Academy of Sciences of the United States of America. 2013;110(39):15740–15745. doi: 10.1073/pnas.1313388110.

[34] Lehto MP, Haag CR. Ecological differentiation between coexisting sexual and asexual strains of *Daphnia pulex*. The Journal of Animal Ecology. 2010;79(6):1241–1250. doi: 10.1111/j.1365-2656.2010.01726.x.

[35] Caron V, Ede FJ, Sunnucks P. Unravelling the paradox of loss of genetic variation during invasion: superclones may explain the success of a clonal invader. PLoS ONE. 2014;9(6). doi: 10.1371/journal.pone.0097744.

[36] Jensen LH, Enghoff H, Frydenberg J, Parker ED. Genetic diversity and the phylogeography of parthenogenesis: comparing bisexual and thelytokous populations of *Nemasoma varicorne* (Diplopoda: Nemasomatidae) in Denmark. Hereditas. 2002;136(3):184–194. doi: 10.1034/j.1601-5223.2002.1360302.x.

[37] Burns M, Hedin M, Tsurusaki N. Population genomics and geographical parthenogenesis in Japanese harvestmen (Opiliones, Sclerosomatidae, *Leiobunum*). Ecology and Evolution. 2018;8(1):36–52. doi: 10.1002/ece3.3605.

[38] Hörandl E. Species concepts in agamic complexes: applications in the *Ranunculus auricomus* complex and general perspectives. Folia Geobotanica. 1998;33(3):335–348. doi: 10.1007/BF03216210.

[39] Kearney M, Blacket MJ, Strasburg JL, Moritz C. Waves of parthenogenesis in the desert: evidence for the parallel loss of sex in a grasshopper and a gecko from Australia. Molecular Ecology. 2006;15(7):1743–1748. doi: 10.1111/j.1365-294X.2006.02898.x.

[40] Burke NW, Bonduriansky R. The geography of sex: sexual conflict, environmental gradients and local loss of sex in facultatively parthenogenetic animals. Philosophical Transactions of the Royal Society B: Biological Sciences. 2018;373(1757):20170422. doi: 10.1098/rstb.2017.0422.

[41] Song Y, Scheu S, Drossel B. Geographic parthenogenesis in a consumer-resource model for sexual reproduction. Journal of Theoretical Biology. 2011;273(1):55–62. doi: 10.1016/j.jtbi.2010.12.020.

[42] Suomalainen E. Parthenogenesis in animals. In: Demerec M, editor. Advances in Genetics. vol. 3. Academic Press; 1950. p. 193–253. Available from: http://www.sciencedirect.com/science/article/pii/S0065266008600863.

[43] Avise JC. Evolutionary perspectives on clonal reproduction in vertebrate animals. Proceedings of the National Academy of Sciences. 2015;112(29):8867–8873. doi: 10.1073/pnas.1501820112.

[44] Allee WCWC. Animal aggregations, a study in general sociology. Chicago: The University of Chicago Press; 1931. Available from: http://archive.org/details/animalaggregatio00alle.

[45] Stephens PA, Sutherland WJ, Freckleton RP. What is the Allee effect? Oikos. 1999;87(1):185–190. doi: 10.2307/3547011.

[46] Møller AP, Legendre S. Allee effect, sexual selection and demographic stochasticity. Oikos. 2001;92(1):27–34. doi: 10.1034/j.1600-0706.2001.920104.x.

[47] R Core Team. R: a language and environment for statistical computing; 2017. Available from: https://www.R-project.org/.

[48] Vrijenhoek RC. Factors affecting clonal diversity and coexistence. American Zoologist. 1979;19(3):787–797.

[49] Pound GE, Cox SJ, Doncaster CP. The accumulation of deleterious mutations within the frozen niche variation hypothesis. Journal of Evolutionary Biology. 2004;17(3):651–662. doi: 10.1111/j.1420-9101.2003.00690.x.

[50] Holt RD, Keitt TH. Alternative causes for range limits: a metapopulation perspective. Ecology Letters. 2000;3(1):41–47. doi: 10.1046/j.1461-0248.2000.00116.x.

[51] Holt RD, Barfield M. Theoretical perspectives on the statics and dynamics of species’ borders in patchy environments. The American naturalist. 2011;178(Suppl 1):S6–25. doi: 10.1086/661784.

[52] Dorken ME, Eckert CG. Severely reduced sexual reproduction in northern populations of a clonal plant, Decodonverticillatus (Lythraceae). Journal of Ecology. 2001;89(3):339–350. doi: 10.1046/j.1365-2745.2001.00558.x.

[53] Courchamp F, Clutton-Brock T, Grenfell B. Inverse density dependence and the Allee effect. Trends in Ecology & Evolution. 1999;14(10):405–410. doi: 10.1016/S0169-5347(99)01683-3.

[54] Wells H, Strauss EG, Rutter MA, Wells PH. Mate location, population growth and species extinction. Biological Conservation. 1998;86(3):317–324. doi: 10.1016/S0006-3207(98)00032-9.

[55] Groom M. Allee effects limit population viability of an annual plant. The American Naturalist. 1998;151(6):487–496. doi: 10.1086/286135.

[56] Lamont BB, Klinkhamer PGL, Witkowski ETF. Population fragmentation may reduce fertility to zero in Banksia goodii – a demonstration of the Allee effect. Oecologia. 1993;94(3):446–450. doi: 10.1007/BF00317122.

[57] Kuussaari M, Saccheri I, Camara M, Hanski I. Allee effect and population dynamics in the Glanville fritillary butterfly. Oikos. 1998;82(2):384–392. doi: 10.2307/3546980.

[58] Perälä T, Kuparinen A. Detection of Allee effects in marine fishes: analytical biases generated by data availability and model selection. Proceedings of the Royal Society B: Biological Sciences. 2017;284(1861):20171284. doi: 10.1098/rspb.2017.1284.

[59] Kaul RB, Kramer AM, Dobbs FC, Drake JM. Experimental demonstration of an Allee effect in microbial populations. Biology Letters. 2016;12(4):20160070. doi: 10.1098/rsbl.2016.0070.

[60] Veit RR, Lewis MA. Dispersal, population growth, and the Allee effect: dynamics of the house finch invasion of Eastern North America. The American Naturalist. 1996;148(2):255–274.

[61] Lewis MA, Kareiva P. Allee dynamics and the spread of invading organisms. Theoretical Population Biology. 1993;43(2):141–158. doi: 10.1006/tpbi.1993.1007.

[62] Roy M, Harding K, Holt RD. Generalizing Levins metapopulation model in explicit space: models of intermediate complexity. Journal of Theoretical Biology. 2008;255(1):152–161. doi: 10.1016/j.jtbi.2008.07.022.

[63] Wang G, Liang XG, Wang FZ. The competitive dynamics of populations subject to an Allee effect. Ecological Modelling. 1999;124(2):183–192. doi: 10.1016/S0304-3800(99)00160-X.

[64] Fowler MS, Ruxton GD. Population dynamic consequences of Allee effects. Journal of Theoretical Biology. 2002;215(1):39–46. doi: 10.1006/jtbi.2001.2486.

[65] Amarasekare P. Allee effects in metapopulation dynamics. The American Naturalist. 1998;152(2):298–302. doi: 10.1086/286169.

[66] Harding KC, Begon M, Eriksson A, Wennberg B. Increased migration in host-pathogen metapopulations can cause host extinction. Journal of Theoretical Biology. 2012;298:1–7. doi: 10.1016/j.jtbi.2011.12.009.

[67] Forsman A. Effects of genotypic and phenotypic variation on establishment are important for conservation, invasion, and infection biology. Proceedings of the National Academy of Sciences. 2014;111(1):302–307. doi: 10.1073/pnas.1317745111.

[68] Agashe D. The stabilizing effect of intraspecific genetic variation on population dynamics in novel and ancestral habitats. The American Naturalist. 2009;174(2):255–267. doi: 10.1086/600085.

[69] Prati D, Peintinger M, Fischer M. Genetic composition, genetic diversity and small-scale environmental variation matter for the experimental reintroduction of a rare plant. Journal of Plant Ecology. 2016;9(6):805–813. doi: 10.1093/jpe/rtv067.

[70] Eckert CG, Samis KE, Lougheed SC. Genetic variation across species’ geographical ranges: the central-marginal hypothesis and beyond. Molecular Ecology. 2008;17(5):1170–1188. doi: 10.1111/j.1365-294X.2007.03659.x.

[71] Orr HA, Unckless RL. Population extinction and the genetics of adaptation. The American Naturalist. 2008;172(2):160–169. doi: 10.1086/589460.

[72] Bell G. Evolutionary rescue and the limits of adaptation. Philosophical Transactions of the Royal Society B: Biological Sciences. 2013;368(1610). doi: 10.1098/rstb.2012.0080.

[73] Uecker H. Evolutionary rescue in randomly mating, selfing, and clonal populations. Evolution. 2017;71(4):845–858. doi: 10.1111/evo.13191.

[74] Johannesson K, Smolarz K, Grahn M, André C. The future of Baltic Sea populations: local extinction or evolutionary rescue? Ambio. 2011;40(2):179–190. doi: 10.1007/s13280-010-0129-x.

